# Dopamine D2 receptors in mossy cells reduce excitatory transmission and are essential for hippocampal function

**DOI:** 10.1101/2023.05.05.539468

**Authors:** Michelle C. Gulfo, Joseph J. Lebowitz, Czarina Ramos, Dong-Woo Hwang, Kaoutsar Nasrallah, Pablo E. Castillo

**Author notes:** To whom correspondence should be addressed: Pablo E. Castillo, MD/PhD Dominick P. Purpura Department of Neuroscience, Albert Einstein College of Medicine 1410 Pelham Parkway South Kennedy Center, Room 703, Bronx, NY 10461, USA, Kaoutsar Nasrallah, PhD, Dominick P. Purpura Department of Neuroscience, Albert Einstein College of Medicine 1410 Pelham Parkway South Kennedy Center, Room 703, Bronx, NY 10461, USA.

## Abstract

Hilar mossy cells (MCs) are principal excitatory neurons of the dentate gyrus (DG) that play critical roles in hippocampal function and have been implicated in brain disorders such as anxiety and epilepsy. However, the mechanisms by which MCs contribute to DG function and disease are poorly understood. Expression from the dopamine D2 receptor (D2R) gene (*Drd2*) promoter is a defining feature of MCs, and previous work indicates a key role for dopaminergic signaling in the DG. Additionally, the involvement of D2R signaling in cognition and neuropsychiatric conditions is well-known. Surprisingly, though, the function of MC D2Rs remain largely unexplored. In this study, we show that selective and conditional removal of *Drd2* from MCs of adult mice impaired spatial memory, promoted anxiety-like behavior and was proconvulsant. To determine the subcellular expression of D2Rs in MCs, we used a D2R knockin mouse which revealed that D2Rs are enriched in the inner molecular layer of the DG, where MCs establish synaptic contacts with granule cells. D2R activation by exogenous and endogenous dopamine reduced MC to dentate granule cells (GC) synaptic transmission, most likely by a presynaptic mechanism. In contrast, removing *Drd2* from MCs had no significant impact on MC excitatory inputs and passive and active properties. Our findings support that MC D2Rs are essential for proper DG function by reducing MC excitatory drive onto GCs. Lastly, impairment of MC D2R signaling could promote anxiety and epilepsy, therefore highlighting a potential therapeutic target.

**SIGNIFICANCE:** Growing evidence indicates that hilar mossy cells (MCs) of the dentate gyrus play critical but incompletely understood roles in memory and brain disorders, including anxiety and epilepsy. Dopamine D2 receptors (D2Rs), implicated in cognition and several psychiatric and neurological disorders, are considered to be characteristically expressed by MCs. Still, the subcellular localization and function of MC D2Rs are largely unknown. We report that removing the *Drd2* gene specifically from MCs of adult mice impaired spatial memory and was anxiogenic and proconvulsant. We also found that D2Rs are enriched where MCs synaptically contact dentate granule cells (GC) and reduce MC-GC transmission. This work uncovered the functional significance of MC D2Rs, thus highlighting their therapeutic potential in D2R- and MC-associated pathologies.

## INTRODUCTION

The human and rodent hippocampus is well-recognized for its roles in spatial learning and memory (1–3). As the main input region to the hippocampus proper that discriminates between sensory inputs, the dentate gyrus (DG) is critical for these functions (1, 2). Hilar mossy cells (MCs) of the DG are crucial yet poorly understood players in hippocampal function (4), including spatial learning and novelty detection, as well as disease processes, such as mood disorders and epilepsy (5–12). MCs have unique anatomical properties that position them to powerfully shape the function of the DG. In addition to mediating local feed-forward inhibition onto dentate granule cells (GCs), each MC sends direct excitatory projections to as many as 35,000 GCs along as much as 75% of the hippocampal axis (13). In turn, GCs send direct excitatory and feed-forward inhibitory projections to MCs locally (14). While MC properties and functions are being elucidated, the molecular and cellular mechanisms underlying MC involvement in critical physiological and pathophysiological processes remain largely unknown.

One hallmark of MCs is their expression from the dopamine D2 receptor gene (*Drd2*) promoter. Remarkably, MCs are the only excitatory hippocampal neurons that express from the *Drd2* promoter in mice (15, 16). Dopamine D2 receptors (D2Rs), along with D3 and D4 receptors, make up the D2-like family of dopamine receptors and are G_i/o_-coupled. Dopamine D1 and D5 receptors comprise the D1-like family of dopamine receptors and are G_s_-coupled D2Rs have been extensively studied throughout the brain for their roles in cognition, mood disorders, schizophrenia, and Parkinson’s disease (17). Quantitative autoradiography for D2Rs in human tissue revealed a marked absence of signal in the GC layer (GCL) and a strong band of signal in the input layers to the GCs (18). In cats, this signal is particularly robust in the inner molecular layer (IML), which mainly contains MC axons targeting GCs (18). Expression from the *Drd2* promoter is a feature of MCs widely used to selectively target them (11, 12, 19–21). One study has tested the role of D2R signaling in MC excitability *in vitro* (22), but the functional significance of MC D2Rs *in vivo* is unknown. Hippocampal dopaminergic signaling has been implicated in processes now associated with MCs, such as spatial memory, novelty detection, anxiety-like behavior, and epilepsy (23–31). In addition, dopamine release in the hippocampus and DG from ventral tegmental area and locus coeruleus fibers (25, 32–34) suggests that MC D2Rs could be activated *in vivo.* Additionally, *Drd2* gene expression is a signature feature of MCs that could be conserved in humans, as supported by D2R autoradiography (18). Therefore, determining the role of MC D2Rs is critical to understanding hippocampal function in health and disease.

To investigate the role of MC D2Rs, we conditionally and selectively removed the *Drd2* gene from MCs in adult mice and assessed the resulting behavioral and cellular phenotypes. We found that this manipulation induced a deficit in spatial memory and promoted anxiety-like behavior, two key modalities regulated by the hippocampus and MCs. In addition, *Drd2* removal from MCs increased the severity of and susceptibility to experimentally-induced seizures. Using a tagged D2R knockin mouse, we revealed that D2Rs are highly expressed in the IML and, consistent with this finding, D2R activation reduced MC-GC synaptic transmission in a presynaptic manner. Thus, our results indicate that MC D2Rs modulate DG functions at least in part by reducing MC-GC transmission.

## RESULTS

### Efficient and selective removal of the *Drd2* gene from hilar mossy cells

To determine the role of MC D2Rs, we selectively removed the *Drd2* gene from MCs. To this end, we bilaterally injected a Cre-expressing virus under the CaMKII promoter (AAV5-CaMKII-mCherry-Cre, or AAV5-CaMKII-mCherry as a control), in the dorsal and ventral hilus of 3-3.5-month-old floxed *Drd2* (*Drd2^fl/fl^*) mice (Fig. 1*A*). As the only excitatory neurons of the hippocampus that express from the *Drd2* promoter are MCs (15, 16), and viral gene expression is under the excitatory *CaMKII* promoter (35), Cre was expected to induce significant loss of *Drd2* expression only in MCs. After 3 weeks were allowed for viral expression, the injection strategy yielded MC *Drd2* conditional knockout (cKO) and Control mice (Fig. 1*A*), and we confirmed this by validating the efficiency, selectivity, and functionality of the viruses. Because of the well-recognized differences in dopamine dynamics and receptor levels between sexes and across the estrous cycle (36, 37), we used male mice as a first approach to test for the functional significance of MC D2Rs.

**Figure 1:**
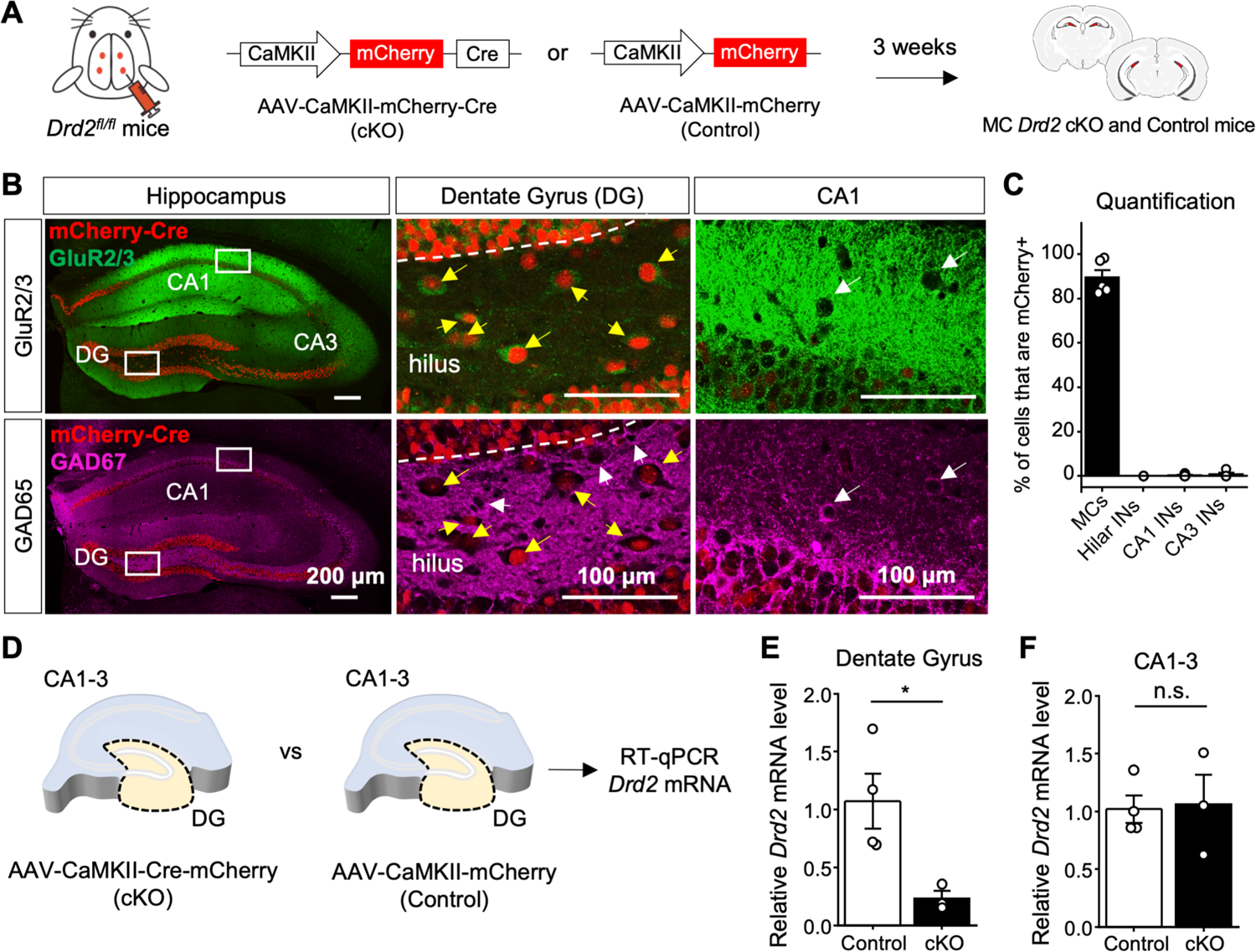
Efficient and selective removal of the *Drd2* gene from hilar MCs. **(A)** Schematic diagram illustrating the experimental strategy to conditionally and selectively KO the *Drd2* gene from MCs in adult mice. AAV5-CaMKII-mCherry-Cre (cKO) or AAV5-CaMKII-mCherry (Control) virus was injected bilaterally into the ventral and dorsal DG of *Drd2^fl/fl^* mice (4 total injection sites) to generate MC *Drd2* cKO or Control mice, respectively. All experiments on these mice were performed at least 3 weeks after viral injections. **(B, C)** Confocal images (B) and quantification (C) revealing high viral expression (mCherry) in hilar MCs (89.4 ± 3.4% of MCs are mCherry positive, N = 5 mice) and a lack of infection in hippocampal interneurons (INs) (0% of hilar INs, 0.3 ± 0.2% of CA1 interneurons and 0.7 ± 0.6% of CA3 interneurons are mCherry positive, N = 5 mice). (B) *Left*, low-magnification images of the hippocampus of a coronal section immunostained for GluR2/3 and GAD67 and assessed for viral efficiency (% of MCs infected) and specificity (% of INs infected). Note that viral expression (mCherry signal) is present throughout the hilus. *Middle,* inset images of the hilus showing that mCherry-expressing hilar neurons (red) are MCs (yellow arrows) identified as GluR2/3-positive (green, top) and GAD67-negative neurons (magenta, bottom). Note the absence of mCherry expression in hilar INs (white arrows, *middle bottom*). *Right,* inset images of CA1 showing examples of mCherry-negative CA1 INs identified as GluR2/3-negative and GAD67-positive (magenta, white arrows). **(D, E)** RT-qPCR analysis. Levels of *Drd2* mRNA relative to *β-actin* mRNA were quantified in the DG and CA1-3 subfields of *Drd2^fl/fl^* mice injected with AAV5-CaMKII-mCherry (Control) *vs* AAV5-CaMKII-mCherry-Cre (cKO) viruses. (D) Schematic showing that DG and CA1-3 sections were dissected from hippocampi of both MC *Drd2* cKO and Control mice for RT-qPCR analysis. (E) Levels of *Drd2* mRNA relative to *β-actin* mRNA in the DG or CA1-3 region for each animal are reported as a fold-difference from that of the Control animals’ mean using the 2^-ΔΔCt^ method (See *SI Appendix*). Relative *Drd2* mRNA was significantly reduced in the DG (Control: 1.1 ± 0.2, N = 4; cKO: 0.2 ± 0.1, N = 3; control vs cKO: p = 0.03, unpaired t-test) but not in CA1-3 areas (control: 1.0 ± 0.1, N = 4; cKO: 1.1 ± 0.3, N = 3; control vs cKO: p = 0.88, unpaired t-test) of cKO mice as compared to Control mice. Data are represented as mean ± SEM. *p < 0.05, n.s. p > 0.5

To test for viral efficiency and selectivity, we performed double immunohistochemistry on injected hippocampal slices for GluR2/3 and GAD67, commonly used in the mouse hilus as markers for MCs and interneurons (INs), respectively (5, 15) (Fig. 1*B*). Quantification of MCs infected with Cre (mCherry positive) confirmed that the virus infected MCs with high efficiency (∼90%) (Fig. 1*C*). Next, we assessed viral selectivity for excitatory neurons to ensure that D2R-expressing INs of CA1-3 (15, 16) were not targeted by Cre. Comparison of the cell types infected with Cre virus (mCherry positive) confirmed that the virus injected in the hilus and driven by the *CaMKII* promoter was indeed selective for excitatory neurons and did not appreciably infect INs of the hilus or CA1 and CA3 regions (Fig. 1*C*). Although mCherry-Cre was expressed in GCs, these excitatory neurons do not express from the *Drd2* promoter (15, 16). We then tested the effectiveness and selectivity of the Cre virus by performing RT-qPCR on injected hippocampal slices for *Drd2* mRNA relative to *β-actin* mRNA. We dissected the DG from the CA regions in hippocampal slices from each animal to separately analyze the two *Drd2*-expressing cell populations of the hippocampus –i.e., MCs of the DG and INs of the CA regions (Fig. 1*D*). We found that the level of *Drd2* mRNA was significantly reduced in the DG of MC *Drd2* cKO animals as compared to Control animals (Fig. 1*E*). In contrast, there was no difference in *Drd2* mRNA level in the CA regions between Control and cKO animals (Fig. 1*F*). These RT-qPCR assessments strongly support that the Cre virus effectively reduced the level of *Drd2* mRNA from Control levels only in MCs of the DG, and not in INs of the CA regions. Having validated our experimental approach, we assessed the behavioral impact of genetic *Drd2* removal from MCs.

#### Deleting the *Drd2* gene from hilar mossy cells impaired object location memory but not object recognition memory

Hippocampal dopaminergic signaling, has been implicated in various forms of spatial memory (23–25, 30). Therefore, we assessed the role of MC D2R signaling in spatial memory by testing MC *Drd2* cKO and Control mice in the Object Location Memory task (OLM). This well-recognized test has been used to study MC function (5, 8, 38). In our experiment, MC *Drd2* cKO and Control mice were allowed to freely explore two identical objects in the training configuration for 4 minutes. One hour later, one object was moved, and mice were allowed to freely explore the two objects in the testing configuration for 5 minutes (Fig. 2*A*). Mice that displayed a preference for the moved object during testing [moved object preference score >55%; moved object preference score = (moved object exploration time/total object exploration time)*100] were considered to pass the test and have intact spatial memory. Mice with a marked preference (>60%) for either object during training and mice with a total exploration under 3 seconds in training or testing were excluded (8, 38). As expected, we found that in training there was no difference in preference for the would-be-moved object between MC *Drd2* cKO and Control mice (Fig. 2*B*). In testing, Control animals passed on average. Their mean moved object preference score was above 55%, and at least 2/3 of animals passed. However, on average, knockout animals failed. Their mean preference score was below 55%, and at least 2/3 of animals failed, indicating a deficit in spatial memory. The difference in performance between cKO and Control animals was significant (Fig. 2*C*). Finally, there was no difference in total object exploration time between Control and cKO animals in training or testing (*SI Appendix*, Fig. S1*A*). In all, these results support that MC D2Rs are essential in spatial memory.

**Figure 2:**
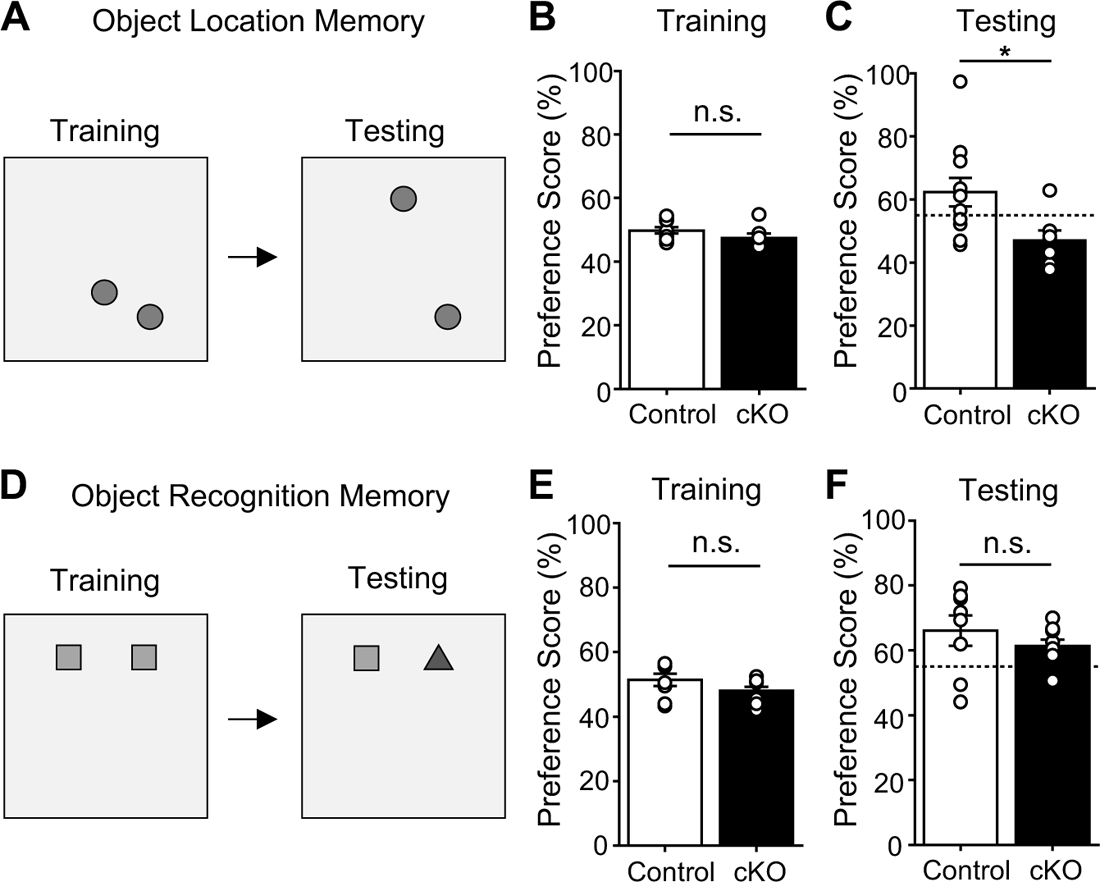
Deleting the *Drd2* gene from hilar MCs impaired OLM but not ORM. **(A-C)** Object Location Memory (OLM) test. (A) OLM test schematic (Training duration: 4 min; Testing duration: 5 min; Interval between training and testing: 1 h). (B) Control and MC *Drd2* cKO mice did not show any significant difference in preference score during training (Control: 49.8 ± 0.9%, N = 11; cKO: 47.5 ± 1.4%, N = 7; Control vs cKO: p = 0.2, unpaired t-test). (C) MC *Drd2* cKO mice showed impaired OLM (mean moved object preference score <55%) with significant reduction in preference for the moved object during testing, as compared to Control mice (Control: 62. 4 ± 4.5%, N = 11; cKO: 47.0 ± 3.1%, N = 7; Control vs cKO: p = 0.025, unpaired t-test). Dashed horizontal line indicate the threshold to pass OLM test, corresponding to a mean preference score > 55%. **(D-F)** Object Recognition Memory (ORM) test. (D) ORM test schematic (Training duration: 4 min; Testing duration: 5 min; Interval between training and testing: 1 h). (E) MC *Drd2* cKO and Control mice did not show significant difference in preference score during training (Control: 51.4 ± 1.9%, N = 8; cKO: 48.0 ± 1.3%, N = 9; Control vs cKO, p = 0.16, unpaired t-test). (F) MC *Drd2* cKO and Control mice both showed no impairment in ORM (mean preference score >55%) and no difference in performance (Control: 66.1 ± 4.6%, N = 8; cKO: 61.3 ± 1.9%, N = 9; Control vs cKO: p = 0.37, unpaired t-test). Dashed horizontal line indicates the threshold to pass the ORM test, corresponding to a mean preference score > 55%. Data are represented as mean ± SEM. *p < 0.05, n.s. p > 0.1.

The role of the hippocampus in recognition memory is more nuanced than its role in spatial memory (39), as other cortical brain regions contribute to this type of memory (38). The DG and MCs have been implicated by several studies in novelty detection and recognition memory (8, 40, 41), although one study reported that inhibiting MC activity has no impact on object recognition memory (5). Nonetheless, novel events trigger dopamine release in the hippocampus (23–25, 34), and particularly hippocampal dopamine D1/D5 receptor signaling has been linked to enhancing object recognition memory (31) and enhancing spatial memory following novelty exposure (23–25). To directly address the potential role of MC D2Rs in recognition memory, we tested MC *Drd2* cKO and Control mice in the Object Recognition Memory task (ORM), which has been used in studies of MC function (5, 8, 38). We used the same scoring scheme and inclusion criteria as for OLM, except that in testing, one object was replaced instead of being moved (Fig. 2*D*). In training, we found no difference in preference for the would-be-replaced object between MC *Drd2* cKO and Control mice (Fig. 2*E*). In testing, both MC *Drd2* cKO and Control animals passed on average. Their mean moved object preference score was above 55%, and at least 2/3 of animals passed. There was no difference in performance between the two groups of mice (Fig. 2*F*). Finally, to probe maximally for a deficit in object recognition memory in MC *Drd2* cKO mice, we challenged the animals to an interval of 24h between training and testing. Still, both groups of mice passed the test, and there was no difference in performance between the two groups (*SI Appendix*, Fig. S1*B*). Our results indicate that MC D2R signaling is not critical for object recognition memory. The results also support that deleting the *Drd2* from MCs does not affect sensory processing and cognition but selectively interferes with spatial memory.

### Deleting *Drd2* from hilar mossy cells promoted anxiety-like behavior

MCs have been implicated in controlling anxiety-like behavior (1, 3, 8–10). The hippocampus can detect conflict and choices to be made between approach and avoidance in the environment, and this detection likely underlies both its roles in spatial memory and anxiety-like behavior (1). To test whether MC *Drd2* cKO mice exhibited alterations in anxiety-like behavior, we first ran them in the Open Field Test (OFT), which assesses locomotor and anxiety-like behaviors (Fig. 3*A*). We found that MC *Drd2* cKO mice spent more time at the edges and avoided the center of the arena as compared to Control mice (Fig. 3*B*), suggesting an increase in anxiety-like behavior (42). The cKO mice also traveled a significantly shorter total distance (*SI Appendix*, Fig. S2*A*) in the center of the Open Field. The reduced time MC *Drd2* cKO mice spent in the center of the arena could not be explained by changes in motor ability including total tracklength and average velocity (Figs. 3*C* and *SI Appendix*, Fig. S2*A*). These results suggest that deleting the *Drd2* gene from MCs promotes anxiety-like behavior. To test this possibility directly, we used the Elevated Plus Maze (EPM) (43). MC *Drd2* cKO and Control mice were allowed to freely explore the EPM for 10 minutes (Fig. 3*D*). We found that MC *Drd2* cKO mice spent significantly less time (Fig. 3*E*) and traveled a significantly shorter total distance (*SI Appendix,* Fig. S2*B*) in the open arms of the maze as compared to Control mice. This anxiety-like phenotype in MC *Drd2* cKO mice could not be explained by motor deficits, as there was no difference in total distance traveled by (Fig 3*F*) or average velocity of (*SI Appendix,* Fig. S2*B*) Control and cKO mice. Altogether, these results indicate that MC D2Rs may have an anxiolytic function.

**Figure 3:**
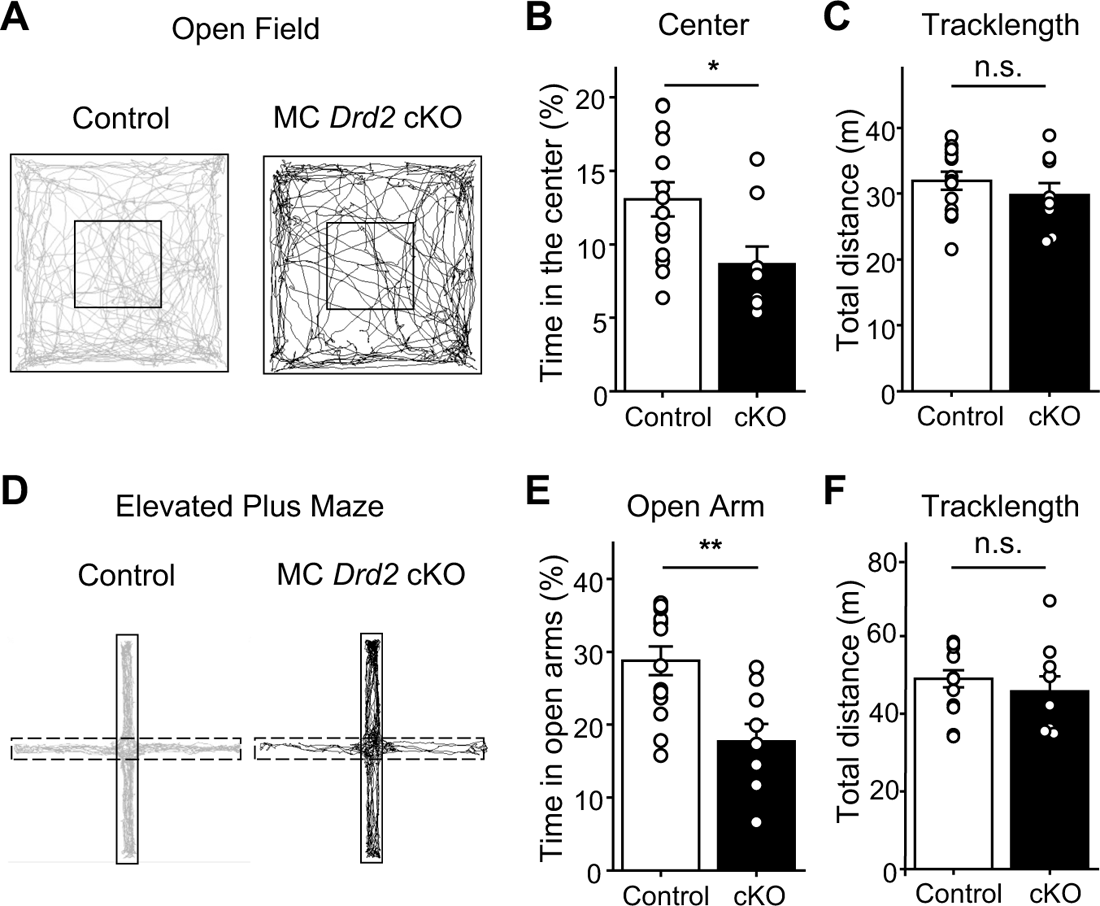
Deleting the *Drd2* gene from MCs promoted anxiety-like behavior. **(A-C)** Open Field Test (OFT). (A) Representative track maps of a Control (*left*, grey trace) and an MC *Drd2* cKO mouse (*right*, black trace) in the open field. (B) MC *Drd2* cKO mice spent a significantly lower percentage of the test time in the center (inner square) of the open field as compared to Control animals (Control: 13.1 ± 1.2%, N = 14; cKO: 8.7 ± 1.2%, N = 9, Control vs cKO: p = 0.014, Mann-Whitney U test). (C) No significant difference in total tracklength was observed between Control and cKO animals (Control: 32.0 ± 1.3%, N = 14; cKO: 29.8 ± 1.8%, N = 9; Control vs cKO: p = 0.34, unpaired t-test). Data are from minutes 0-6 of the test. **(D-F)** Elevated Plus Maze (EPM). (D) Representative track maps of a Control (*left*, gray trace) and an MC *Drd2* cKO mouse (*right*, black trace) in the EPM. Dashed lines correspond to the open arm. (E) MC *Drd2* cKO mice spent a significantly lower percentage of test time in the open arms of the EPM than Controls (Control: 28.8 ± 2.0%, N = 14; cKO: 17.7 ± 2.4%, N = 9, Control vs cKO: p = 0.0019, unpaired t-test). No significant difference in total tracklength was observed between Control and cKO animals (Control: 49.0 ± 2.2 m, N = 14; cKO: 45.8 ± 3.9 m, N = 9, Control vs cKO: p = 0.45, unpaired t-test). Data are from minutes 0-10 of the test. Data are represented as mean ± SEM. n.s. p > 0.2, *p < 0.05, **p < 0.01.

### Deleting the *Drd2* gene from hilar mossy cells increased seizure severity and susceptibility

In addition to anxiety-like behavior, MCs play a key role in temporal lobe epilepsy (4, 5, 11, 12). There is also evidence that dopamine regulates seizures from the limbic system (28). Germline deletion of *Drd2* is pro-convulsive and excitotoxic, particularly in the CA3 area (27, 29). Pharmacologic studies also support an anti-epileptic role of D2R signaling in the hippocampus (28). Therefore, we examined the contribution of MC D2Rs in regulating seizure activity using the well-established kainic acid model of acute seizure induction (44) (Fig. 4*A*). MC *Drd2* cKO and Control mice were injected with kainic acid (20 mg/kg, i.p.) and their seizure stage was scored every 10 minutes for 2 hours according to the modified Racine scale (*SI Appendix*). MC *Drd2* cKO mice had a higher cumulative Racine score than Control mice across the scoring period, with an approximately 2-fold difference present by the end of it (Fig. 4*B*). Thus, over the 2-hour scoring period, MC *Drd2* cKO mice had significantly more severe seizures than Control mice (Fig. 4*B,C*). The cKOs also had a greater susceptibility to seizures, as they reached the convulsive seizure stage significantly sooner than Controls did (Fig. 4*D*). These results support that MC D2R signaling can act as a powerful negative regulator of seizure activity.

**Figure 4:**
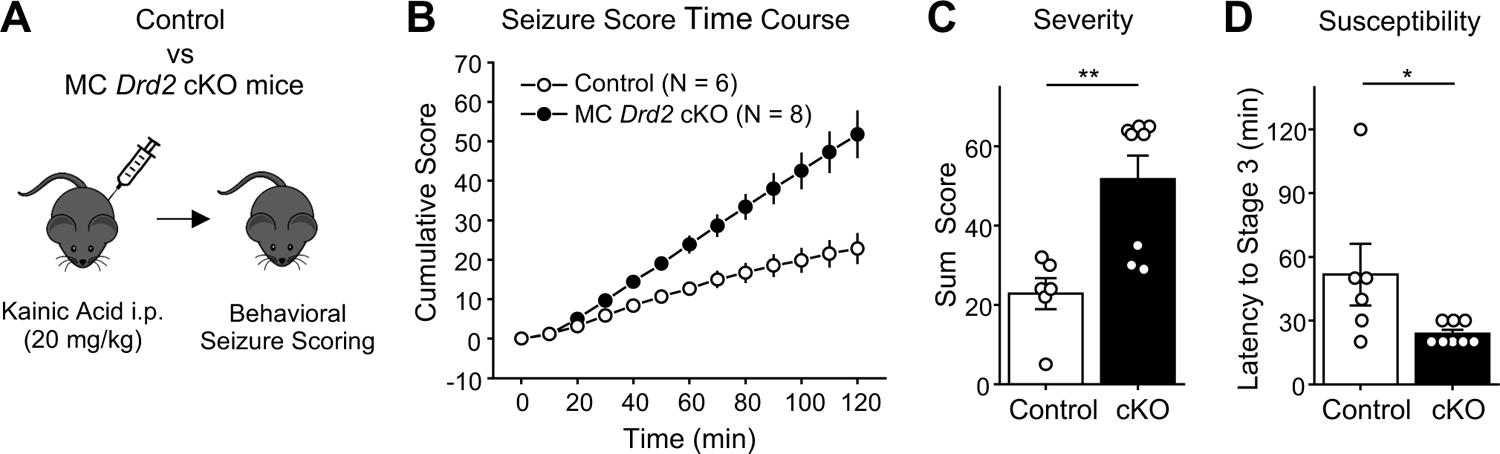
Deleting the *Drd2* gene from MCs increased KA-induced seizure severity and susceptibility. **(A)** Experimental timeline. Seizures were acutely induced using a single KA i.p. injection (20 mg/kg) and scored using a modified Racine scale for 120 min. **(B-D)** The cumulative Racine score at the end of 120 minutes of scoring (B) was significantly greater in MC *Drd2* cKO mice than Control mice (C, severity; Control: 22.8 ± 3.9%, N = 6; cKO: 51.8 ± 6.0%, N = 8; Control vs cKO: p = 0.0095, Mann-Whitney U test). MC *Drd2* cKO mice also showed a significant decrease in latency to convulsive seizures as compared to Control mice (D, susceptibility; Control: 51.7 ± 14.5%, N = 6; cKO: 23.8 ± 1.8%, N = 8; Control vs cKO: p = 0.025, Mann-Whitney U test). Data are presented as mean ± SEM. *p < 0.05; **p < 0.01.

### D2Rs are enriched in the inner molecular layer of the dentate gyrus

Having uncovered a role for MC D2Rs *in* vivo, we sought to determine potential cellular and molecular mechanisms underlying the phernotypes we observed. We began by investigating the subcellular distribution of D2Rs in MCs. Given the low specificity and sensitivity of D2R antibodies for addressing such question in tissue (17), we used a knockin mouse with a superecliptic pHluorin (SEP) epitope fused to the N-terminus of endogenous D2Rs (45, 46). Live labeling of SEP-D2Rs was achieved by incubating *ex vivo* coronal slices containing the DG with an α-GFP antibody prior to permeabilization. At low power, SEP-D2R signal appeared as a distinct band surrounding the GCL as visualized with DAPI, corresponding to the IML (Fig. 5*A*). This IML signal was not present when the coronal slice was not incubated with the α-GFP antibody (Fig. 5*B*). When imaged at 63X using AiryScan, SEP-D2R signal bounding the GCL appeared as puncta (Fig. 5*C*), similar to the punctate distribution of D2Rs previously observed in the midbrain (45, 46). Intensity analysis of SEP-D2R signal confirmed the clear enrichment of D2Rs in the IML relative to that measured in the GCL and MML (Fig. 5*D*). The lack of expression from the *Drd2* promoter in GCs (15, 16) strongly suggests that the IML signal arises from MC axons. Notably, no SEP-D2R puncta were observed on the MC cell bodies in the hilus which were identified by GluR2/3 labeling (Fig. 5E). The enrichment of D2Rs in the IML with no apparent surface receptors on the somatodendritic compartment of MCs, supports the hypothesis that D2R acts to decrease transmitter release from MC axon terminals.

**Figure 5:**
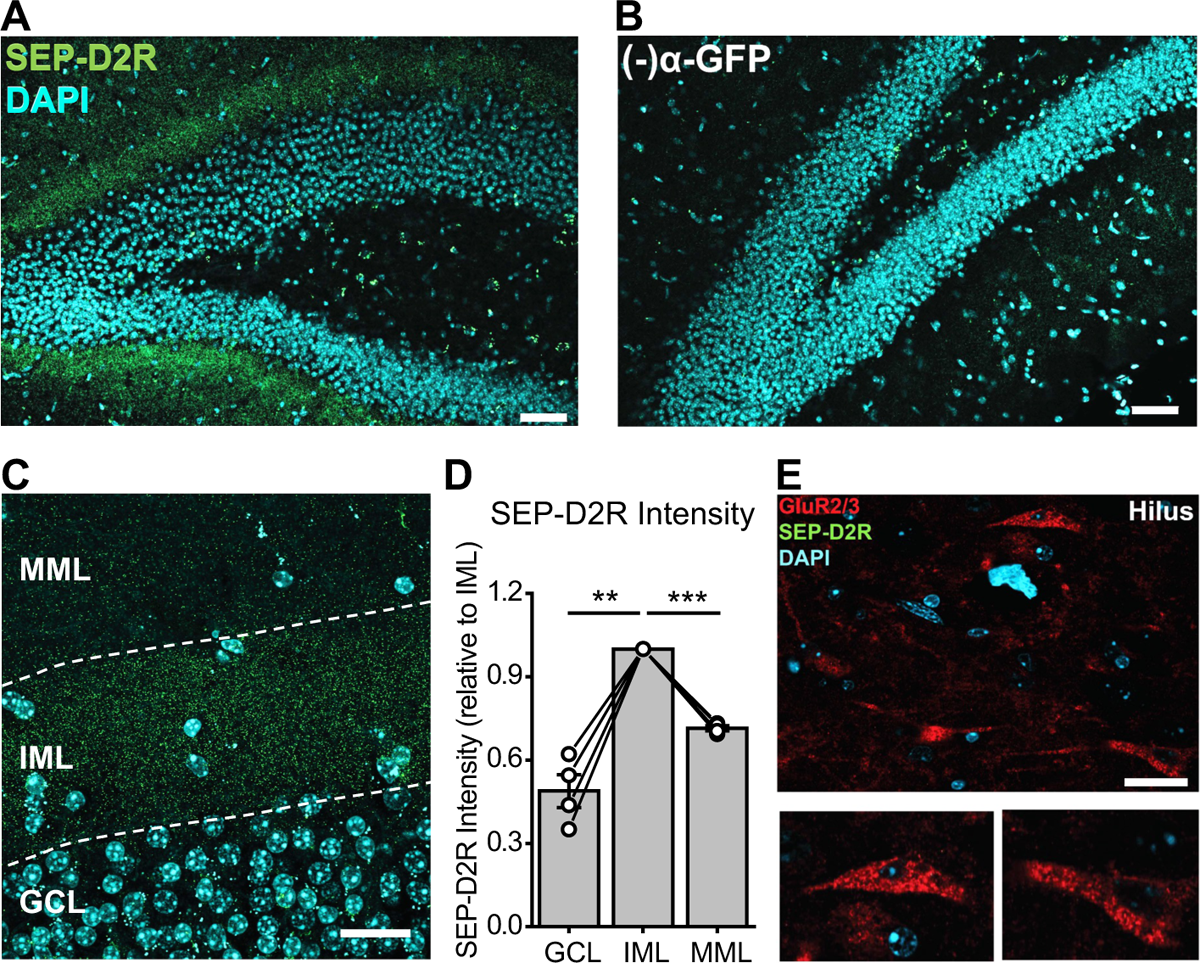
Enriched D2R expression in the inner molecular layer of the dentate gyrus. **(A)** 20X image of D2R labeling in the DG in a SEP-D2R knockin mouse following signal amplification with an α-GFP antibody conjugated to AlexFluor488. The SEP epitope was live-labeled prior to permeabilization to selectively visualize surface D2Rs. Scale: 100 µm. **(B)** 20X image of the DG in an SEP-D2R mouse that was not incubated with the α-GFP antibody. Imaging and visualization parameters are identical to those used in A. Scale: 100 µm. **(C)** Maximum intensity projection of a z-stack (thickness = 2.94 µm) taken at 60X of SEP-D2R labeling in the IML. SEP-D2R labeling is abundant in the IML where MC terminals reside with less apparent labeling in the GCL and MML. Scale: 20 µm. **(D)** Intensity quantification of SEP-D2R signal following amplification in the GCL, IML, and MML imaged in single z-planes at 60X. (IML vs. GCL: p = 0.0055; IML vs MML: p = 0.0001; n = 4 images/2 slices/2 animals; Dunnett’s multiple comparison test following repeated-measures one-way ANOVA). **(E)** 60X image of GluR2/3 labeling in the hilus following antibody-amplification of SEP-D2R. Surface SEP-D2R puncta are absent from GluR2/3 containing cell bodies (insets). Data are represented as mean ± SEM. ** p < 0.01, *** p < 0.001

### D2R activation depresses mossy cell-granule cell excitatory transmission by a presynaptic mechanism

Our results thus far supported that MC D2Rs, likely expressed in MC axon terminals, are critical to hippocampal function *in vivo*. We therefore hypothesized that a potential mechanism by which MC D2R signaling mediates its effects on behavior is by modulating MC-GC synaptic transmission, which we tested using hippocampal slice electrophysiology. We performed whole-cell voltage-clamp recordings of GCs and elicited MC-GC excitatory postsynaptic currents (EPSCs) by electrically stimulating MC axons in the IML with inhibitory synaptic transmission blocked –i.e., in the presence of the GABA_A_ receptor antagonist picrotoxin (100 µM) and the GABA_B_ receptor antagonist CGP55845 (3 µM). We found that bath application of dopamine (20 µM, 15 min) significantly and reversibly depressed MC-GC EPSCs (Fig. 6*A,B*). This depression was blocked in the presence of the competitive D2-like antagonist sulpiride (1 µM) (Fig. 6*A,B*). In addition, it was accompanied by an increase in paired-pulse ratio (PPR) (Fig. 6*C*), consistent with a presynaptic mechanism. D2R antagonism had no effect on baseline transmission (Fig. 6*D*), suggesting that MC D2Rs are not tonically active at the MC-GC synapse. The dopamine-mediated depression of MC-GC synaptic transmission was not accompanied by changes in GC holding current or input resistance (Fig. 6 *E*), also consistent with a presynaptic mechanism. Deleting *Drd2* from MCs had no significant effect on basal neurotransmitter release, as indicated by the lack of PPR change (Fig. 6*F*), further supporting the absence of tonic activity of MC D2Rs on MC-GC transmission. In contrast, MC *Drd2* removal precluded the dopamine-mediated depression of MC-GC transmission (Fig. 6*G*), supporting that dopamine depresses MC-GC transmission by targeting presynaptic D2Rs. Lastly, we examined whether endogenous dopamine could also modulate MC-GC synaptic transmission. To test this possibility, we use amphetamine, which potently releases dopamine from dopaminergic terminals (47). Bath application of D-amphetamine hemisulphate (20 µM) also induced a reversible reduction of MC-GC transmission that was abolished in the continuous presence of sulpiride (1 µM) (Figure 6*H*). Together, these findings indicate that activation of MC D2R by exogenous and endogenous dopamine reversibly reduced MC-GC transmission, most likely by inhibiting glutamate release from MC axon terminals.

**Figure 6:**
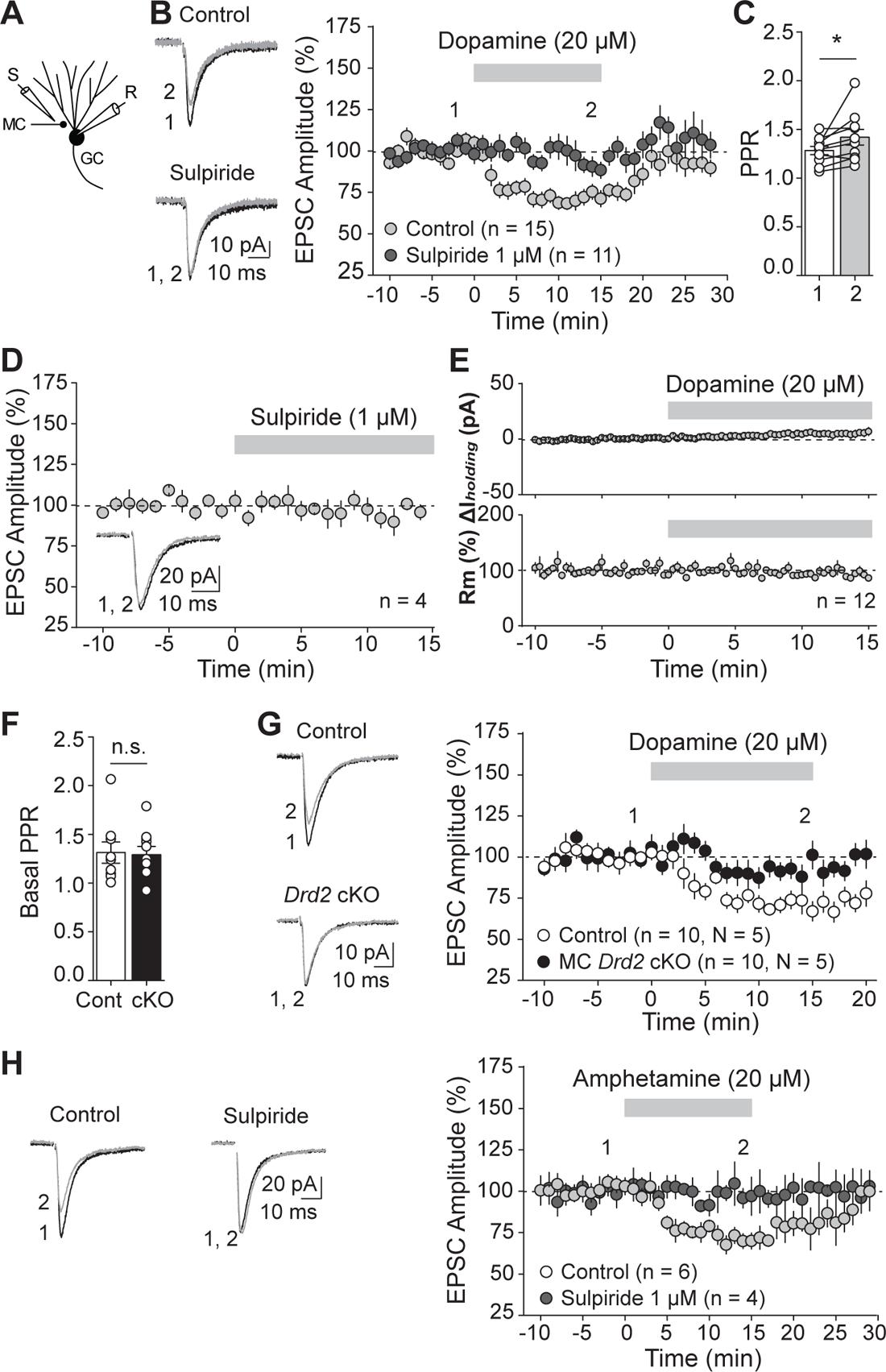
Activation of MC D2Rs depressed MC-GC excitatory transmission. **(A-E)** Dopamine depressed MC-GC synaptic transmission in a D2-like receptor-dependent manner. (A) Recording configuration for MC-GC transmission. (B) Representative average traces (*left*) and time-course summary plot (*right*) showing that bath application of dopamine (20 µM, 15 min) reduced MC-GC transmission in a reversible manner in control wild-type animals (ontrol: 71.3 ± 3.4% of baseline, n = 15, p < 0.0001, paired t-test). This reduction was blocked in the continuous presence of the D2-like receptor antagonist sulpiride (1 µM) (sulpiride: 93.5 ± 5.9% of baseline, n = 11, p = 0.1, paired Wilcoxon signed ranks test; control vs sulpiride: p = 0.0009, Mann-Whitney U test). The reduction was also associated with significant increase in PPR (C, pre: 1.28 ± 0.43, dopamine: 1.42 ± 0.08, n = 6, pre vs dopamine: p = 0.048, paired t-test) (D) Representative average traces and time-course summary plot showing that bath application of sulpiride (1 µM, 15 min) did not significantly affect MC-GC EPSC amplitude (95.0 ± 5.0% of baseline, n = 4, p = 0.34, paired t-test). (E) No significant changes in GC holding current (ΔI_holding_, top: 5.4 ± 3.2 pA difference from baseline, n = 12, p = 0.19, paired t-test) or GC membrane resistance (Rm, *bottom*: 96.5 ± 5.2% of baseline, n = 12, p = 0.60, paired t-test) were observed during dopamine application in B. **(F,G)** Dopamine depressed MC-GC transmission via MC D2Rs. (F) Basal PPR was similar in Control and MC *Drd2* cKO animals (Control: 1.31 ± 0.11%, n = 9; cKO: 1.29 ± 0.08%, n = 10; Control vs cKO: p = 0.97, Mann-Whitney U test). (G) Representative average traces (*left*) and time-course summary plot (*right*) showing that dopamine-mediated depression of MC-GC transmission was abolished in MC *Drd2* cKO animals as compared to Control mice (Control: 70.7 ± 4.1% of baseline, n = 10, p = 0.00025, paired t-test; cKO: 93.5 ± 4.3% of baseline, n = 10, p = 0.41, paired Wilcoxon signed ranks test; Control vs cKO: p = 0.0007, Mann-Whitney U test). **(H)** Amphetamine depressed MC-GC transmission in a D2-like receptor-dependent manner. Representative average traces (*left*) and summary plot (*right*) showing that bath applying D-amphetamine hemisulphate (20 µM, 15 min) reversibly depressed MC-GC synaptic transmission in control (70.8 ± 2.9% of baseline, n = 6, p = 0.0019, paired t-test) but not in the continuous presence of sulpiride (1 µM) (sulpiride: 99.2 ± 8.7% of baseline, n = 4, p = 0.99, paired t-test; control vs sulpiride, p = 0.041, unpaired t-test). Data are represented as mean ± SEM. *p < 0.05, n.s. p > 0.3.

Although a presynaptic mechanism of dopamine-mediated depression of MC-GC transmission is consistent with the D2R enrichment in the IML where MCs synaptically contact GCs (Fig. 5), D2R expression in other MC compartments and synaptic inputs cannot be discarded, especially given the low sensitivity of the SEP-D2R KI approach (45). However, we found that dopamine application had no significant effect on MC active and passive properties (e.g., rheobase, number of action potentials per injected current step, and input resistance –see Methods) monitored under pharmacological blockade of excitatory and inhibitory synaptic transmission (*SI Appendix,* Fig. S3*A-E*). In addition, dopamine did not affect spontaneous EPSC amplitude and frequency in MCs (*SI Appendix,* Fig. S3*F-G*) or evoked GC-MC EPSCs (*SI Appendix*, Fig. S3*H*). Thus, these results support our anatomical and functional findings (Figs. 5 and 6), indicating that MC D2Rs may primarily modulate MC and DG functions by reducing glutamate release from MC projections to GCs.

## DISCUSSION

This study reveals a role of MC D2R signaling in crucial aspects of hippocampus-dependent cognitive function and diseases. By selectively and conditionally removing D2Rs from MCs in adult mice, we demonstrate that MC D2Rs are essential for spatial memory and play anxiolytic and anticonvulsant roles. At the cellular level, dopamine negatively controlled MC-GC synaptic transmission via MC D2R activation, while it did not significantly impact MC excitability or excitatory inputs. Furthermore, we show anatomical evidence for D2R enrichment in MC axons. In light of the extensive and powerful connections that MCs make to GCs, it follows that MCs strongly regulate DG function. Given the diffuse nature of dopamine release and extensive dopaminergic projections within the hippocampus, dopaminergic signaling via presynaptic D2Rs emerges as an ideally suited neuromodulatory mechanism for controlling MC-GC excitatory transmission and hippocampal function. Our study provides mechanistic insights into MC and D2R function in memory, anxiety-like behaviors, and seizures, suggesting MC D2Rs as a potential therapeutic target.

It is well-established that hippocampal dopaminergic signaling is required for hippocampus-dependent learning and memory (34, 48). Novel events trigger dopamine release to the hippocampus (23–25, 34), and pharmacologic and knockout studies support a role for both hippocampal D1-like and D2-like receptors in both spatial and object recognition memory (48, 49). We gathered functional and anatomical evidence supporting that MC D2Rs negatively regulate the excitatory output of MCs onto GCs via a presynaptic mechanism. Previous autoradiography work suggested that D2Rs are expressed in MC axons while absent from GCs in cats and humans (18). Presumably due to the low specificity and sensitivity of D2R antibodies, several studies have inferred the expression of dopamine receptors from mRNA analysis (17). In the DG, the expression of a reporter from the *Drd2* promoter occurred in MCs but not GCs and INs (15, 16). Taking advantage of the SEP-tagged D2R KI mice (45, 46), we discovered enriched D2R expression in putative MC axons. In contrast, we did not detect D2Rs in MC somatodendritic compartment. Our functional analyses using the endogenous ligand dopamine did not detect any D2R-dependent modulation of MC active or passive membrane properties (*SI Appendix*, Fig. S3). Intriguingly, a previous study reported that the D2-like agonist quinpirole increases MC excitability (22), but whether a selective D2R antagonist can block this effect is unclear. While our results discarded that D2Rs regulate MC functional properties and excitatory inputs onto MCs (*SI Appendix*, Fig. S3), we cannot exclude additional effects at MC-INs synapses. However, the proconvulsant effect observed in MC *Drd2* cKO mice (Fig. 4) suggests that MC D2R signaling has a net inhibitory role in the DG.

Consistent with a presynaptic localization of D2Rs, we found that their activation with dopamine triggered a significant and reversible reduction in MC-GC transmission, which was associated with PPR increase and abolished by D2-like antagonism and *Drd2* deletion from MCs. The D2R-mediated suppression of glutamate release is likely due to the G_i/o_-mediated inhibition of presynaptic calcium influx via voltage-gated calcium channels and/or activation of potassium channels (17). In addition, the psychostimulant amphetamine, known to potently promote release of dopamine, also mediated a reversible D2R-dependent depression of MC-GC synaptic transmission. A previous study reported that transient bath application of dopamine and amphetamine induced a D2R-dependent, presynaptic long-term depression at neighboring perforant path inputs onto GCs (50). While the mechanism by which D2R activation induces this plasticity remains unclear, distinct D2R downstream signaling could account for the different duration of the D2R-mediated depression across synapses.

MC-GC excitatory transmission can powerfully activate GCs throughout the DG (51). We have recently reported that retrograde endocannabinoid signaling strongly suppresses glutamate release from MC axon terminals by activating presynaptic Type 1 cannabinoid receptors, another G_i/o_-coupled receptor highly expressed in MC axon boutons (52). Endocannabinoid signaling is typically induced by GC activity, thereby suppressing MC inputs onto active GCs only. In contrast, the extensive dopaminergic projection throughout the hippocampus (3, 25) strongly suggests that dopamine effectively inhibits MC excitatory drive onto GCs by diffusely targeting MC D2Rs. Thus, dopaminergic and endocannabinoid signaling may have distinct but complementary ways of controlling MC-GC synaptic transmission and DG network activity. Such synergism mediated by different presynaptic G_i/o_-coupled receptors likely represents a general motif throughout the brain that regulates neuronal communication and behavior (53).

Deleting *Drd2* from hilar MCs selectively impaired OLM but not ORM. These findings are consistent with recent studies implicating MCs in spatial memory (5–7). For instance, optogenetic inhibition of MCs impaired OLM but not ORM (5). Similarly, inhibiting MCs by overexpressing Kir2.1 potassium channels interfered with spatial memory retrieval but did not affect ORM (6). It is well-established that hippocampus-dependent learning and memory requires normal hippocampal dopaminergic signaling (34, 48). Pharmacological and germline knockout studies support a role for hippocampal D2-like receptors in both spatial and object recognition memory (34, 48, 49). However, our study is the first to directly address the cell-specific function of MC D2Rs in DG-dependent behaviors. The DG plays a critical role in spatial memory discrimination (54–56), a computational process that minimalizes the overlap between similar neural representations, thereby reducing memory interference. GC sparse activity, which is classically attributed to their unique intrinsic properties and their particularly strong inhibition, is believed to be critical for spatial pattern separation (57). Remarkably, exposure to novel objects triggers dopamine release in the hippocampus (23–25, 34). By reducing MC excitatory drive onto GC (Fig. 6), MC D2Rs engaged during the OLM could improve signal to noise and enable memory discrimination.

Our findings supporting an anxiolytic function of MC D2Rs are consistent with previous pharmacological studies reporting that D2R signaling can have anxiolytic effects *in vivo* and suggest that D2Rs on MCs significantly contribute to this effect. For instance, D2R agonists subcutaneously administered decreased mouse open-field thigmotaxis (42) and reduced rat ultrasonic vocalizations (58). Our data also provide a potential mechanism for the role of MCs in anxiety-like behaviors. Three independent studies reported that MC activity is anxiolytic in the EPM (8–10), although others did not observed this action (5, 6). Chemogenetic and optogenetic activation of MCs increased mouse open-arm time (8, 10), and specific removal of MCs by diphtheria toxin reduced mouse open-arm time in the EPM (9). Anxiety tests often involve a spatial component, so lack of orientation might yield equal preference for anxiogenic and non-anxiogenic environments. MC *Drd2* cKO mice do not have equal preference for the open and closed arms of the EPM but rather spend ∼80% of their time in the closed arms (Fig. 3). Similarly, they spent ∼90% of their time at the edges of the open field. The behavior of MC *Drd2* cKO mice in the EPM (Fig. 3) and in response to kainic acid (Fig. 4) suggest an anxiolytic and net inhibitory role of MC D2Rs in the DG. It is worth noting that anxiety can interfere with animals’ exploration of novel objects or environments, which is critical for memory task performance (1). However, we observed no significant difference in total exploration time between objects when comparing MC *Drd2* cKO vs control animals. The spatial memory deficit and anxiety-like behavior observed in MC *Drd2* cKO mice likely reflect a state of disordered DG information processing, possibly related to the decision of approach and avoidance, which has been proposed previously to underlie both phenotypes (1, 4).

Recent studies showed that chemogenetic inhibition of MCs reduces experimentally-induced seizures by kainic acid and pilocarpine (11, 12). Further, MC-GC synapses are robustly strengthened following initial seizures, whereas reducing MC-GC transmission reduces seizure activity (11, 12, 59). Our findings demonstrating that MC D2Rs depress MC-GC transmission and play an anticonvulsant role are consistent with these findings. However, in the chronic mouse model of temporal lobe epilepsy, activation of surviving MCs is antiepileptic (5, 60), suggesting that the function of MC D2Rs may also differ significantly with the disease stage. There is good evidence that D2Rs play an antiepileptic role (28). For example, germline D2R knockout animals have a substantially higher seizure score than wild-type animals (29). In addition, D2R antagonists used as antipsychotics promote seizures in epileptic and nonepileptic patients, and anti-parkinsonian treatments that stimulate D2Rs are antiepileptic (28). D2R expression is reduced in the temporal lobe of epileptic patients and in rodent models of temporal lobe epilepsy (28). Given the diffuse nature of dopamine release and extensive dopaminergic projections in the hippocampus (3, 25), signaling through MC D2Rs may be a mechanistic explanation for the antiepileptic role of D2Rs. Moreover, by dampening MC-GC transmission, MC D2Rs emerge as a potential target to dampen seizure activity.

In conclusion, our study supports that MC D2Rs serve an inhibitory role that facilitates information processing in the DG, spatial memory task performance, and normal exploration of anxiogenic environments. In addition, MC D2Rs likely prevent DG runaway activity that occurs during epileptic seizures, and they may also be implicated in schizophrenia, cognitive and mood disorders. Finally, strong D2R expression in the IML is conserved across species perhaps as a testament to the importance of MC D2Rs in hippocampal function including in humans. Tools to selectively activate MC D2Rs *in vivo* will be required to demonstrate the therapeutic potential of these receptors.

## MATERIAL AND METHODS

*Drd2*^fl/fl^ (B6.129S4(FVB)-Drd2tm1.1Mrub/J, Strain #: 020631; The Jackson Laboratory), C57BL/6 (C57BL/6NCrl, Strain Code: 027; Charles River Laboratories), and SEP-D2R knockin mice (B6;129S7-*Drd2^tm1.1Jtw^*/J, Strain #: 030537; The Jackson Laboratory) were used in this study. All animals were group housed in a standard 12:12 h light:dark cycle and had free access to food and water. Animals were bred, cared for, handled, and used according to protocols approved by the Institutional Animal Care and Use Committee (IACUC) at Albert Einstein College of Medicine and the Vollum Institute (OHSU), in accordance with guidelines from the National Institutes of Health (NIH). Experimental procedures for MC *Drd2* conditional KO generation, confirmation of AAV expression and immunolabeling, examination of *Drd2* mRNA expression by RT-qPCR, behavioral testing (OF, ORM, OLM and EPM), seizure induction and monitoring, visualization of hippocampal D2Rs using SEP-D2 knockin mice, acute hippocampal slice preparation, electrophysiology, and biocytin visualization for *post hoc* confirmation of MC identity are detailed in SI Appendix, Supplementary Materials and Methods. Details of image acquisition and quantification, and of all statistical analyses performed are also included in SI Appendix, Supplementary Materials and Methods. For all additional details, refer to SI Appendix, Supplementary Materials and Methods

## ACKNOWLEDGMENTS

We thank all members of the Castillo lab for constructive feedback. We thank Maria Gulinello, director of the Albert Einstein College of Medicine Animal Behavior Core, for training for and supervision of animal behavior experiments, Aubrey Siebels for assistance with analysis of MC spontaneous activity, and Subrina Persaud for assistance with immunostaining, imaging, and brain and tissue processing in early stages of this project. We also thank Miwako Yamasaki (Hokkaido University) and Teresa A. Milner (Weill Cornell Medicine) for their attempt to detect D2Rs in the hippocampus using immunohistochemistry and immunoelectron microscopy, respectively. Finally, special thanks to John T. Williams (Vollum Institute, OHSU) for permitting Joseph J. Lebowitz to perform the experiments using SEP-D2R knockin mice. This research was supported by the National Institutes of Health (NIH), R01-NS113600, R01-MH125772; R01-NS115543 and R01-MH116673 to P.E.C.; F31-MH122134 to M.C.G.; and F31-MH127810 to C.Z.. M.C.G. was partially supported by T32-GM007288; J.J.L. was supported by R01-DA004523 and T32-DA007262 grants; and DWH was supported by R35-GM136296. Confocal images were obtained at the Einstein Imaging Core (supported by The Rose F Kennedy Intellectual Disabilities Research Center - shared instrument grant NIH 1S10OD25295 to Konstantin Dobrenis).

## Author contributions

M.C.G., K.N. and P.E.C. conceptualized, designed research and wrote the manuscript; M.C.G. performed and analyzed all behavioral experiments, brain slice electrophysiology, and immunostainings; K.N. performed and analyzed the experiments involving experimental seizures and some electrophysiology experiments, as well as performed cell quantification and RT-qPCR slice preparation; J.J.L. performed and analyzed the experiments using SEP-D2R knockin mice, C.R. performed and analyzed experiments to assess MC intrinsic properties; D.W.H. performed and analyzed RT-qPCR experiments. All authors read and edited the manuscript.

## Data Availability

All study data are included in the article and/or *SI Appendix*. This study did not generate new unique reagents. Further information and requests for resources and reagents should be directed to and will be fulfilled by the lead contact, Pablo E. Castillo. Any additional information required to reanalyze the data reported in this paper is available from the lead contact upon request.

The authors declare no competing interests.

## SUPPORTING INFORMATION (SI Appendix)

### SUPPLEMENTARY MATERIALS AND METHODS

#### Experimental model and subject details

*Drd2^fl/fl^* (B6.129S4(FVB)-Drd2tm1.1Mrub/J, Strain #: 020631; The Jackson Laboratory), C57BL/6 (C57BL/6NCrl, Strain Code: 027; Charles River Laboratories) and SEP-tagged D2R KI mice (B6;129S7-*Drd2^tm1.1Jtw^*/J, Strain #: 030537, The Jackson Laboratory) were used in this study. Exon 2 of the *Drd2* gene is flanked with loxP sites in *Drd2^fl/fl^*mice. All animals were housed in groups of 2-5 siblings of the same sex in individually ventilated cages (IVC) with access to food and water *ad libitum*, on a standard 12-hr light/12-hr dark cycle. Animals were bred, cared for, handled, and used according to protocols approved by the Institutional Animal Care and Use Committee (IACUC) at Albert Einstein College of Medicine and the Vollum Institute (OHSU), in accordance with guidelines from the National Institutes of Health (NIH).

#### MC *Drd2* conditional KO

*Drd2^fl/fl^* male mice (12-14 weeks old for immunolabeling, RT-qPCR, and behavioral experiments including seizure induction, and 5-7 weeks old for electrophysiology experiments) were anesthetized with isoflurane (up to 5% for induction and 1–2.5% for maintenance) and placed in a stereotaxic frame (Kopf Model 940). Cre-expressing virus (AAV5-CaMKII-mCherry-Cre, 5.8 x10^12^ virus molecules/mL, UNC Vector Core), or Control virus (AAV5-CaMKII-mCherry, 4.9 x10^12^ virus molecules/mL, UNC Vector Core) was bilaterally injected (0.5 μL/site, at 0.1 μL/min) into both the dorsal (1.9 mm posterior, 1.2 mm lateral, 2.2 mm ventral) and ventral (3.2 mm posterior, 2.2 mm lateral, 2.8 mm ventral) hilus to generate MC *Drd2* cKO or Control mice, respectively. Immunolabeling, RT-qPCR, behavioral experiments, and electrophysiology experiments were performed 3 to 6 weeks post-injection.

#### Confirmation of AAV expression and immunolabeling

For every animal injected for MC *Drd2* cKO and Control experiments, expression of the AAV reporter mCherry was assessed bilaterally in dorsal, medial, and ventral DGs. Animals were included in the study if mCherry signal was present throughout the hilus in at least four of the six sites examined, thus if at least approximately two-thirds of all MCs in the brain were infected with virus. For RT-qPCR and electrophysiology, we evaluated 400 and 300 µm-thick slices, respectively, floating in the chamber of on an upright electrophysiology microscope (Nikon Eclipse FN1) with fluorescence attachments. These slices were prepared as described in the sections below, “Examination of *Drd2* mRNA expression by RT-qPCR” and “Acute hippocampal slice preparation for electrophysiology.” For behavior experiments (including seizure induction), we evaluated 50 µm-thick fixed sections mounted on slides (ProLong Diamond Antifade Mountant with DAPI, ThermoFisher) under an upright fluorescent microscope (Zeiss) to confirm viral expression *post hoc*. These sections were prepared in the same manner as sections were for immunolabeling assessments of viral efficiency and specificity (Fig. 1B*,C*), described as follows.

Injected mice were deeply anesthetized with isoflurane (3-5%) and transcardially perfused with a solution containing 4% paraformaldehyde (PFA) in 1X phosphate buffer saline (PBS). Brains were extracted and fixed in 4% PFA for 24-48 hours at 4°C. After fixation, brains were stored in 1X PBS at 4°C, and 24-48 hours later, 50 µm-thick coronal floating sections were prepared in 1X PBS from the brains using a vibratome (VT1000 S, Leica Microsystems Co.). After collection, slices were stored at −20°C in a cryoprotectant solution (30% ethylene glycol anhydrate, 30% glycerol, 25% 1X PBS, 15% ddH2O, % = vol/vol*100). A minimum of one day following section preparation, sections were washed in 1X PBS, 3 x 10 min each, at room temperature (RT). They were then incubated in blocking solution (10% goat serum, 1% Triton X-100 in 1X PBS) for 2 hours at RT. Primary antibody solution (5% goat serum, 0.1% Triton X-100, and primary antibodies in 1X PBS) was then applied for 48 hours at 4°C. Primary antibodies to label MCs (Rabbit anti-GluR2/3, 1:50; EMD Millipore/Chemicon, Cat# AB1506) and INs (Mouse anti-GAD67, 1:500; EMDMillipore/Chemicon, Cat# MAB5406) were used. Sections were then washed in 1X PBS, 4 x 10 min each, at RT. Secondary antibodies to visualize the primary antibodies for MCs and INs were used (Alexa Fluor 488 Goat anti-Rabbit, 1:500; Invitrogen, Cat# A11008; Alexa Fluor 647 Goat anti-Mouse, 1:1000; Invitrogen, Cat# A32728). Sections were then washed in 1X PBS, 4 x 10 min each, at RT. The second to last of these washes included 40,6-diamidino-2-phenylindole (DAPI, 1:1000 in PBS, 20 min, ThermoFisher) to label cell nuclei. Finally, sections were mounted onto microscope slides using ProLong Diamond Antifade Mountant (ThermoFisher) and allowed to dry in the dark before confocal imaging.

#### Image acquisition and cell quantification

Coronal hippocampus-containing sections stained for GluR2/3 and GAD67 (see above) were imaged by confocal microscopy. Tile scan images were acquired using the 10X objective (1.5 zoom) of a Zeiss LSM 880 Airyscan Confocal microscope with SuperResolution, and ZEN (black edition) software. For viral efficiency and specificity quantification, images were analyzed in Fiji and cells were counted using the Cell Counter Fiji plug-in by a blind experimenter. All cells were confirmed for total counts by the presence of nuclear staining in the DAPI channel. Viral efficiency was calculated as percentage of MCs infected with virus [(Number of hilar neurons positive for mCherry, GluR2/3, and DAPI, and negative for GAD67) / (Number of hilar neurons positive for GluR2/3 and DAPI, and negative for GAD67)]*100. Viral specificity was calculated as % of INs in the hilus, CA1, and CA3 infected with virus: [(Number of neurons positive for mCherry, GAD67, and DAPI, and negative for GluR2/3) / (Number of neurons positive for GAD67 and DAPI, and negative for GluR2/3)]*100. Per animal, three hippocampi (one dorsal, medial, and ventral) with hiluses that would be considered infected with virus for the animal inclusion criteria described above were analyzed. Average efficiency and specificity values across the three hippocampi were averaged per animal and reported (Fig. 1C). Figure 1 includes representative images and quantification of viral efficiency and specificity. Right and left hippocampus were randomly used.

#### Examination of *Drd2* mRNA expression by RT-qPCR

Quantitative reverse transcription PCR (RT-qPCR) was used to assess expression of the *Drd2* mRNA, in DG vs CA1-3 of both control and MC *Drd2* cKO mice. Acute transversal hippocampal slices were prepared from these mice according to the methods described in “Acute hippocampal slice preparation for electrophysiology” with the following modifications. Before slicing, tools, equipment, and surface areas were cleaned with RNase*Zap* (Invitrogen AM9780) to eliminate RNases. In addition, 400-µm thick slices were cut and incubated for 30 min in artificial cerebrospinal fluid (ACSF, composition below) solution at RT before processing. After this incubation, 12 hippocampi per animal (4 dorsal, 4 medial, and 4 ventral) were selected for RNA isolation and analysis. Each hippocampus was dissected into DG and CA1-3 portions, and all portions per animal were placed in 2 separate tubes containing TRIzol® Reagent (Thermo Fisher, 15596026) to isolate total RNA from both regions per animal and were frozen. As there were 7 animals total (4 Control, 3 cKO) and one DG and one CA1-3 region sample per animal, 14 total samples were obtained.

Total RNA (315 ng) was converted to cDNA by a reverse transcriptase enzymatic reaction with SuperScript® III First-Strand Synthesis System (Thermo Fisher, 18080051). Quantitative PCR reaction was prepared by mixing 15 ng of cDNA per reaction, 4 μM of primer pair (i.e. forward and reverse) and Power SYBR Green PCR Master Mix (Thermo Fisher, 4358577). PCR runs were performed on the ViiA 7 Real-Time PCR system (Thermo Fisher). Each reaction for individual primer pairs was composed of 3 technical replicates. Normalization of the *Drd2* expression within each of the 14 samples was performed by subtracting the average Ct value (i.e. cycle number where the exponential amplification of cDNA molecules occurs) of endogenous control gene (*β-actin* mRNA, *Actb*) from the average Ct value of the *Drd2* mRNA (i.e. ΔCt). In summary, a ΔCt value was calculated for each sample as that sample’s *Drd2* Ct –*Actb* Ct. Subsequently, the mean ΔCt value of all Control DG samples and the mean ΔCt value of all control CA region samples were used as reference points to compare relative expression of the *Drd2* mRNA across all samples of the DG or CA1-3 regions, respectively. A ΔΔCt value was calculated for each sample as Sample ΔCt – Control Mean ΔCt. The ΔΔCt values from each sample were then converted to a fold-difference, calculated as 2^-ΔΔCt^ for each sample. These fold-differences reflect the relative *Drd2* expression within and across samples, for each sample. The 2^-ΔΔCt^ values from each sample (14 total: one DG and one CA1-3 sample per MC *Drd2* cKO and Control animal) were plotted, grouped according to hippocampal region and condition, using OriginPro software (OriginLab 2015, 2023). The sequence information for the primers utilized in RT-qPCR can be found in the table below.

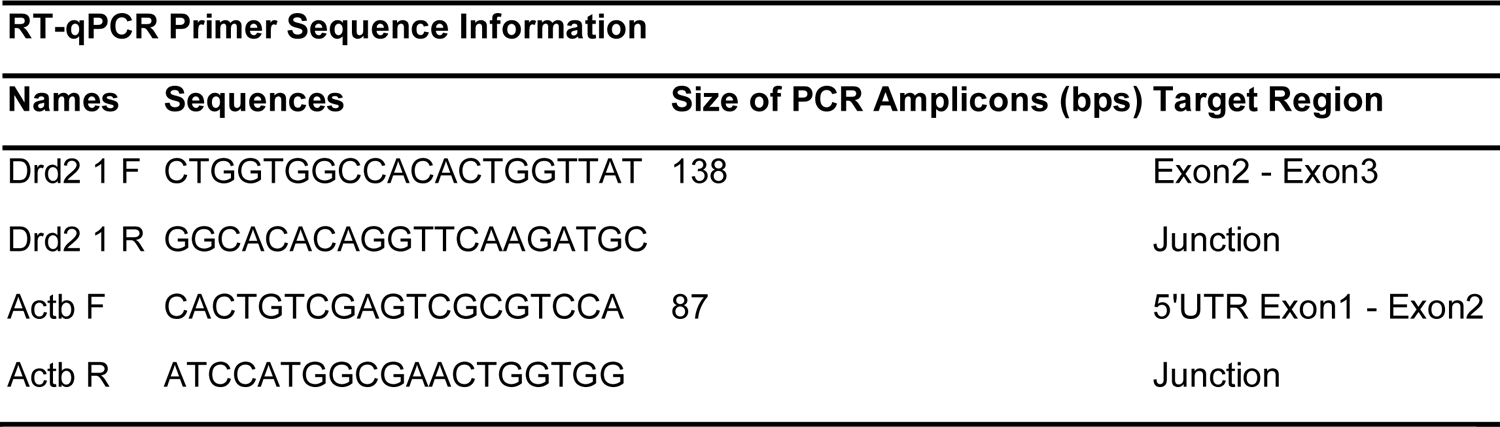

#### Behavioral testing

Behavioral analysis was performed on 3.75-5-month-old *Drd2^fl/fl^* injected males, starting 3 weeks post viral injection. Mice of this age have a sufficiently high and stable cognitive functioning for behavioral assessment (1). All behavioral tests were conducted and scored by an experimenter blind to condition (Control vs cKO). Viral expression was verified *post hoc* in each animal in fixed slices (see “Confirmation of AAV expression and immunolabeling”). Tests were completed between 8 AM – 4 PM in dedicated procedure rooms, in dim lighting and with soft background noise. At the end of each session, mice were returned to their home cage and left undisturbed until the next test. All mice were habituated and run in all tests individually. The arena and maze were cleaned with 70% ethanol between mice in both training and testing. Viewer software, with tracking based on the mouse’s body, was used to record and analyze mouse behavior.

#### Open Field Test (OFT)

Each mouse was run in the OFT (Fig. 3A*-C*) 8 min after the 4 min handling session. In this test, each mouse was allowed to freely explore the open field (square arena: 40.5 cm length x 40.5 cm width x 30.5 cm height) for 9 minutes. The bottom of the open field was a layer of white matte shelf liner, and the top was open. The arena walls were rigid (acrylic) and lined with the same shelf liner, and each wall contained one black visual cue that would all remain for subsequent memory tests. All surfaces were waterproof and glare-free. Measurements of tracklength, velocity, and center duration were calculated in 3 bins of 3 minutes. The center was defined in the software as an 18.4 x 18.4 cm square in the center of the arena. As tracking was performed based on the mouse’s body, brief nose or head entrances into the center were not counted as part of the center duration or tracklength. The arena was cleaned with 70% ethanol between mice.

#### Object Location Memory (OLM)

The OLM test (Fig. 2A*-C*) was run in the OF arena. During training, mice were allowed to freely explore two identical, visually rich objects, for 4 minutes. 1 hour after training, each mouse was allowed to freely explore the two objects, for 5 minutes, in the testing configuration where one of the objects was moved 12 cm from its original position. The identity of the moved object was randomized. The experimenter live-scored the time animals spent exploring each object during training and testing. Preference score for the moved object was calculated as [(moved object exploration time)/(moved + unmoved object exploration time)]*100. Mice that displayed a preference for the moved object during testing (moved object preference score >55%) were considered to pass the OLM test and have intact spatial memory. Preference score during training was calculated in the same manner but for the object that would be moved during testing. Consistent with previously employed criteria (2, 3), mice that had a marked preference (>60%) for either object during training or that had a total object exploration time less than 3 seconds in training or testing were excluded. The arena was cleaned with 70% ethanol between mice in both training and testing.

#### Object Recognition Memory (ORM)

The ORM test (Fig. 2D*-F*) was also run in the OF arena. During training, mice were allowed to freely explore two identical objects for 4 minutes. For testing, one of the objects was replaced with another novel object (new object) but kept in the same position as the original object. The other object (old object) was not replaced or moved. One hour after training, the testing session was performed in which each mouse was allowed to freely explore the old and new objects for 5 minutes. Two pairs of novel objects (not used in OLM) were used in this test to enable counterbalancing of which object was novel during testing within each cohort. Which position was novel (left or right) and which object pair was used first in counterbalancing was randomized between cohorts. The experimenter live scored the time the animals spent exploring each object during training and testing. Preference score for the new object was calculated as [(new object exploration time)/(new + old object exploration time)]*100. Mice that displayed a preference for the new object during testing (new object preference score >55%) were considered to pass the test and have intact object recognition memory. Preference score during training was calculated in the same manner but for the object that would be replaced during testing. As in the OLM test, mice had a marked preference (>60%) for either object during training or that had a total object exploration time less than 3 s in training or testing were excluded. A second ORM test was run with a 24 h retention interval, or time between training and testing, to probe maximally for ORM deficits in MC *Drd2* cKO mice (Fig. S1*B*). The same procedures were followed as in the 1 h retention interval ORM test described above except for the following: training and testing duration were 10 and 5 minutes, respectively, and two new pairs of objects were used.

#### Elevated Plus Maze (EPM)

The EPM is a plus-shaped maze with 4 equally-sized rectangular arms (50 x 10 cm) united by a square-shaped center (10 x 10 cm), elevated 50 cm above the floor. It consists of two closed arms enclosed on three sides by opaque walls (40 cm high), and two open arms. For testing in the EPM (Fig. 3D*-F*), each animal was placed in the center and allowed to freely explore the maze for 10 minutes. Measurements of tracklengths, velocity, and duration in open and closed arms were calculated in 2 bins of 5 minutes. The open and closed arm zones were defined in the software as beginning where a mouse’s body could be fully in a given arm. All EPM testing was completed by 2 PM.

#### Seizure induction and monitoring

To induce seizures, MC *Drd2* cKO and Control mice were injected with 20 mg/kg kainic acid (KA, HelloBio HB0355) prepared in 0.9% NaCl the same day, via intraperitoneal route (i.p.). Mice were monitored and behavioral seizures were scored for 120 minutes post-KA injection by an experimenter blind to condition (Control vs cKO) according to a modified Racine scale (4). The scale is defined as stage 0: normal behavior, stage 1: immobility and rigidity, stage 2: head bobbing, stage 3: forelimb clonus and rearing, stage 4: continuous rearing and falling, stage 5: clonic-tonic seizure, stage 6: death. The maximum Racine score was recorded every 10 minutes and the cumulative seizure score was obtained by summing these scores across all 12 bins of the 120 min period.

#### Visualization of dentate gyrus D2Rs using SEP-D2 knockin mice

SEP-D2R knockin mice and live-labeling strategies to visualize surface D2Rs have been previously characterized (5, 6). For visualizing surface D2Rs in the DG, *ex vivo* slices in the coronal plane were generated from male SEP-D2R mice at 2-4.5 months of age. Krebs solution used for brain extraction and cutting was warmed to 32-35°C and contained (in mM): 126 NaCl, 2.5 KCl, 1.2 MgCl_2_, 2.4 CaCl_2,_ 1.4 NaH_2_PO_4_, 25 NaHCO_3_, and 11 dextrose. Slices were allowed to recover at 34°C for 30 min prior to live-labeling. Solutions used for cutting and recovery both contained 10 µM MK-801 and were continuously bubbled with 95% O_2_ / 5% CO_2_. For live labeling, slices were incubated in bubbled Krebs solution with the inclusion of an α-GFP primary antibody conjugated to AlexaFluor 488 (Invitrogen, cat #: A-21311) at 1:400 for 1 hour at 34°C. Slices were washed by incubation in fresh Krebs for 15 min at 34°C following labeling. Fixation was achieved with 4% PFA for 30 min at RT. Slices were blocked and permeabilized for 1 hour at RT in PBS containing 0.5% Triton X-100 and 10% normal goat serum (Southern Biotech, cat#: 0060-01). To label GluR2/3, slices were incubated for 48 hours in 0.1% Triton X-100 and 5% normal goat serum with α-GluR2/3 (Millipore, cat#: AB1506) at 1:50. Secondary antibody for GluR2/3 was Goat-anti-Rabbit AlexaFluor-555 (Invitrogen, cat#: A-21429) and was incubated for 1 hour at RT at 1:500 in 0.1% Triton X-100 and 5% normal goat serum. Slices were washed 3x in PBS for ≥ 20 min following primary and secondary incubation. Slices were mounted on glass slides with Fluoromount-G with DAPI (Southern Biotech, cat#: 0100-20).

Imaging was carried out on a Zeiss LSM-980 at the OHSU Advanced Light Microscopy Core. For low power imaging, 20X objective was used with a 75 µm pinhole to maximize signal. Control tissue not exposed to α-GFP antibody was imaged on the same day using identical imaging parameters. High-power images were obtained using Zeiss Airyscan with a 63X objective and a zoom of 1.7. For z-stacks, slices were taken with a step size of interval of 0.140 µm and the image displayed as a maximum intensity projection. Intensity measurements were made using Zeiss Zen imaging software by drawing ROI’s around the GCL, IML, and MML and obtaining the average fluorescent intensity for SEP-D2R signal. All image processing for final presentation was conducted in Zen imaging software.

#### Acute hippocampal slice preparation for electrophysiology

Electrophysiology experiments were performed on acute transversal hippocampal slices prepared from 8-12-week-old mice. C57BL/6 “wild-type” mice were used for all experiments, except in Fig. 6F*,G* where Control and MC *Drd2* cKO mice were used. Animals were anesthetized with isoflurane and euthanized in accordance with Einstein IACUC regulations. After decapitation, the brain was removed rapidly from the skull and immediately placed in an ice-cold N-methyl-D-glucamine (NMDG) solution equilibrated with 95%O_2_ / 5% CO_2_. NMDG solution contained (in mM): 93 NMDG, 2.5 KCl, 1.25 NaH_2_PO_4_, 30 NaHCO_3_, 20 HEPES, 25 D-glucose, 2 Thiourea, 5 Na-Ascorbate, 3 Na-Pyruvate, 0.5 CaCl_2_, 10 MgCl_2_, and 93 HCl. The solution was pH-adjusted to 7.35 using 1M HCl. The left and right hippocampi were carefully dissected from the brain, secured in a 4% agar mold, and cut into 300-400 µm-thick slices in the NMDG solution using a vibratome (VT1200 S Microslicer, Leica Microsystems Co.). Slices collected off the vibratome were transferred to a chamber containing an extracellular ACSF solution equilibrated with 95% O_2_ / 5% CO_2_. The ACSF solution contained (in mM): 124 NaCl, 2.5 KCl, 26 NaHCO_3_, 1 NaH_2_PO_4_, 2.5 CaCl_2_, 1.3 MgSO_4_, and 10 D-glucose. Its pH was 7.4. The chamber which contained the slices was kept in a 34°C water bath, and 15 minutes after the last slice was collected the chamber was moved to RT. Slices were allowed to recover at RT for at least 40 minutes before recording. Slices across the hippocampal dorsoventral axis that contained all structures/lamellae were used for recording.

### Electrophysiology

For all recordings, slices were held in a submersion-type recording chamber kept at 28±1°C and perfused with ACSF at 2 mL/min. Whole-cell patch-clamp recordings from dentate GCs or MCs were performed using a Multiclamp 700A amplifier (Molecular Devices) in voltage clamp or current clamp mode. Neurons were voltage clamped at −60 or −70 mV as described (V_h_ = −60 or −70 mV). The patch-type pipette electrodes (∼3-4 MΩ) contained a K^+^-based internal solution of the following composition (in mM): 135 KMeSO_4_, 5 KCl, 1 CaCl_2_, 5 NaOH, 10 HEPES, 5 MgATP, 0.4 Na_3_GTP, 5 EGTA and 10 D-glucose. The solution was adjusted to pH 7.25 and 287-293 mOsm. Throughout the recordings, series resistance was monitored with a −5 mV, 80 ms voltage step. Series resistance across cells ranged from ∼10-30 MΩ for GCs and ∼17-21 MΩ for MCs. Any cell that exhibited a significant change in series resistance (> 20%) across the experiment was excluded from analysis.

Mature GCs were targeted for recording by blind patching in the upper blade of the granule cell layer. The lower half of the granule cell layer near the hilus was avoided as this zone has a higher proportion of adult-born GCs. Mature GCs were identified by characteristic hyperpolarized resting membrane potential (RMP, checked immediately after membrane break in, −72 to −83 mV). Cells with high membrane resistance (> 800 MΩ) were considered as putative adult born GCs (7) and were excluded from the analysis. MC-evoked EPSCs were recorded in GCs (Fig. 6), in the continuous presence of GABA_A_ receptor antagonist picrotoxin (100 µM) and GABA_B_ receptor antagonist CGP55845 (3 µM) to block inhibitory transmission. To activate MC axons, a stimulating patch-type micropipette filled with ACSF was broken slightly at the tip (∼10–20 μm) and placed in the IML close (< 40 μm) to the border of the granule cell layer. To elicit synaptic responses in GCs, two monopolar square-wave voltage pulses (100 μs pulse width, 5-30 V, 100 ms inter-stimulus interval) were delivered through a stimulus isolator (Digitimer DS2A-MKII) every 20 seconds. Stimulation intensity was adjusted to obtain EPSCs of comparable magnitude (30-80 pA) across experiments.

Putative MCs were patched visually in the hilus. MC identity was confirmed by the presence of a characteristic high frequency of spontaneous EPSCs (in Fig S3*F-H*, when excitatory synaptic transmission was intact), by checking the firing pattern in response to depolarizing step of currents (non-burst firing and action potentials with almost no afterhyperpolarization) (8) (Fig. S3*A-H*), and by using *post hoc* morphological analysis (Fig. S3*A-E*). In experiments evaluating MC intrinsic properties (Fig. S4*A-E*), excitatory and inhibitory synaptic transmission was blocked with NBQX (10 µM), D-APV (50 µM), picrotoxin (100 µM) and CGP55845 (3 µM). MCs were voltage-clamped at −70 mV (V_h_ = −70 mV) to prevent non-experimental firing. RMP was checked immediately after break-in, and several sweeps in voltage clamp were acquired to calculate membrane input resistance. To determine rheobase and action potential frequency across a range of injected current, 0 to +400 pA of current (500 ms duration) was injected in 40 steps of +10 pA every 10 seconds. In Fig. S3*F-H*, spontaneous and evoked EPSCs were recorded from MCs (V_h_ = −60 mV) in the continuous presence of picrotoxin (100 µM) and CGP55845 (3 µM). In Fig S3*H*, GC axons were locally stimulated with a bipolar theta glass pipette filled with ACSF (∼10–20 μm tip) placed in the DG subgranular zone to evoke EPSCs. To elicit synaptic responses in MCs, two monopolar square-wave voltage pulses (100 μs pulse width, 5-70 V, 100 ms inter-stimulus interval) were delivered through a stimulus isolator (Digitimer DS2A-MKII) every 20 seconds. Stimulation intensity was adjusted to obtain EPSCs of comparable magnitude (30-100 pA) across experiments. The Group II mGluR agonist DCG-IV (1 μM), which selectively reduces GC synaptic transmission (9, 10), was applied at the end of each experiment to confirm that EPSCs arose from GC stimulation.

Reagents were bath-applied following dilution into ACSF from stock solutions stored at −20°C prepared in water or DMSO, depending on the manufacturer’s recommendation. D-Amphetamine hemisulphate and Dopamine-HCl were prepared fresh daily by suspending powder stored at RT (for D-Amphetamine) or 4°C (for Dopamine-HCl) in H_2_O. These fresh stock solutions were kept on ice in the dark and used within 3 hours.

Electrophysiological data were acquired at 5 kHz, filtered at 2.4 kHz, and analyzed using a Multiclamp 700A amplifier (Molecular Devices) and custom-made software for IgorPro 7.01 (Wavemetrics Inc.). PPR was defined as the ratio of the amplitude of the second EPSC (baseline taken 1-2 ms before the stimulus artifact) to the amplitude of the first EPSC. Thirty consecutive sweeps were averaged to calculate PPR (Fig. 6). The magnitudes of dopamine- and amphetamine-mediated depressions, and the lack of effect of sulpiride, were determined by comparing responses from the final 5 min baseline with responses 10-15 min after wash-in (Fig. 6). Averaged traces include 20 consecutive individual responses. For each MC recorded to assess intrinsic properties, membrane input resistance values from 10 sweeps acquired in voltage clamp were averaged (Fig S3*E*), the numbers of action potentials resulting from each current step were counted manually (Fig. S3*C*), and rheobase was reported as the smallest of the 40 current steps that induced at least one action potential (Fig. S3*D*). Spontaneous EPSC events were analyzed using the Igor software Neuromatic Plugin. For each recording, average event amplitude and inter event interval were calculated from events that occurred during the final 20 sweeps of baseline and dopamine incubation (defined above) (Fig S3*G*). Representative traces (Fig S3*F*) were selected from the final 10 consecutive sweeps of baseline and dopamine incubation.

#### Biocytin visualization for *post hoc* confirmation of MC identity

The identities of putative MCs patched in the presence of NBQX (Fig. S3*A-E*) was confirmed anatomically *post hoc* using biocytin, as NBQX can alter MC firing properties (13). MC were defined by the presence of thorny excrescences (11) and a triangular soma with one major dendritic branch arising from all three poles (12). Biocytin (0.2%) was included in the internal solution used to patch MCs. After the electrophysiology recordings, acute hippocampal slices were fixed in 4% PFA for 24 hours at 4°C. They were then washed in 1X PBS for 20 min followed by 1X + 0.2% TritonX-100, 3 x 15 min each, at RT, Next, they were labelled with an Alexa Fluor 594-conjugated streptavidin-containing solution (Alexa Fluor 594-conjugated streptavidin, ThermoFisher, diluted 1:400 in 1X PBS + 0.2% Triton X-100) overnight at RT. Finally, they were washed in 1X PBS, 3 x 5 min each, at RT and mounted using ProLong Diamond Antifade Mountant with DAPI (ThermoFisher). Images were acquired using the 60X oil-immersion objective of a Zeiss LSM 880 Airyscan Confocal microscope with SuperResolution, and ZEN (black edition) software.

#### Quantification and Statistical Analysis

When testing for potential differences between two populations of data, both distributions were first assessed for normality using the Shapiro-Wilk test. If both distributions were normal, student’s two-tailed t-tests were used. Paired t-tests were used for within-group comparisons and unpaired two-sample t-tests were used for between-group comparisons. For non-normal distributions, non-parametric tests were used. Paired sample Wilcoxon signed rank and Mann-Whitney U tests were used respectively for within-group and between-group comparisons. SEP-D2R intensity values relative to the IML were analyzed by repeated-measures one-way ANOVA followed by Dunnett’s multiple comparison tests to directly assess differences between groups.

Statistical significance was set to p < 0.05 (*** indicates p < 0.001, ** indicates p < 0.01, * indicates p < 0.05, and n.s. indicates not significant with p > 0.05). All data are represented as mean ± SEM. N refers to number of animals and n refers to number of replicates per animal. Unless otherwise stated, all experiments included at least three animals per condition. SEP-D2R intensity values were analyzed in Prism (GraphPad Prism Software 9.4.1) and all other statistical analyses were performed using OriginPro Software (OriginLab 2015, 2023). All graphs were made using OriginPro Software.

---

**Figure S1.**
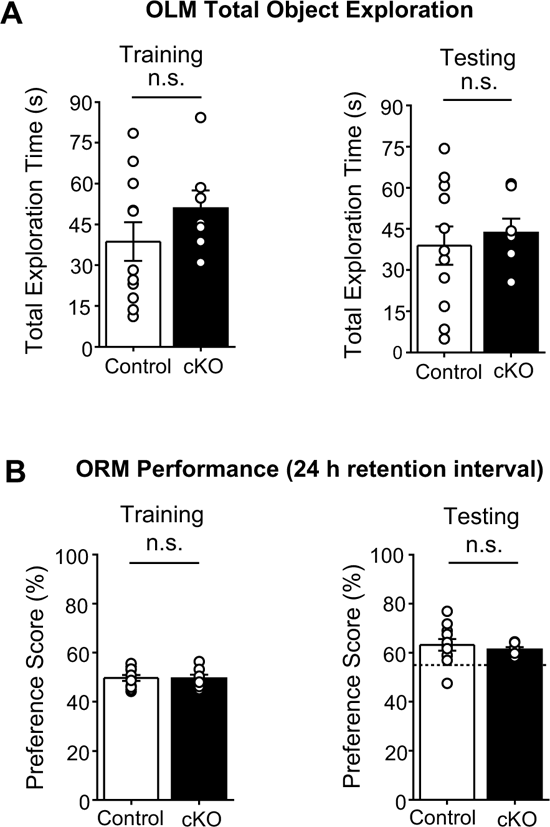
Total object exploration in the OLM test and performance in the 24h-retentional interval ORM test by MC *Drd2* cKO and Control mice. **(A)** Total object exploration in the Object Location Memory (OLM) test. There was no significant difference in total object exploration time between Control and MC *Drd2* cKO mice during training (*left,* Control: 38.7 ± 7.10 s, N = 11; cKO: 50.9 ± 6.58 s, N = 7; Control vs cKO: p = 0.254, unpaired t-test) or testing (*right*, Control: 38.9 ± 7.02 s, N = 11; cKO: 43.6 ± 5.03 s, N = 7; Control vs cKO: p =0.633, unpaired t-test). **(B)** Object Recognition Memory (ORM) test with a 24 h retention interval (Training duration: 10 min; Testing duration: 5 min; Interval between training and testing: 24 h). *Left*, Control and MC *Drd2* cKO mice showed no difference in preference score during training (Control: 49.7 ± 1.21%, N = 11; cKO: 49.5 ± 1.53%, N = 7; Control vs cKO: p = 0.936, unpaired t-test). *Right,* Control and MC *Drd2* cKO mice showed intact ORM (mean preference score >55%), and no difference in performance (Control: 63.3 ± 2.41%, N = 11; cKO: 61.4 ± 0.880, N = 7; Control vs cKO: p = 0.487, unpaired t-test). Dashed horizontal line indicates the threshold to pass ORM test, corresponding to a mean preference score >55%. Data are represented as mean ± SEM. n.s. p > 0.2

**Figure S2.**
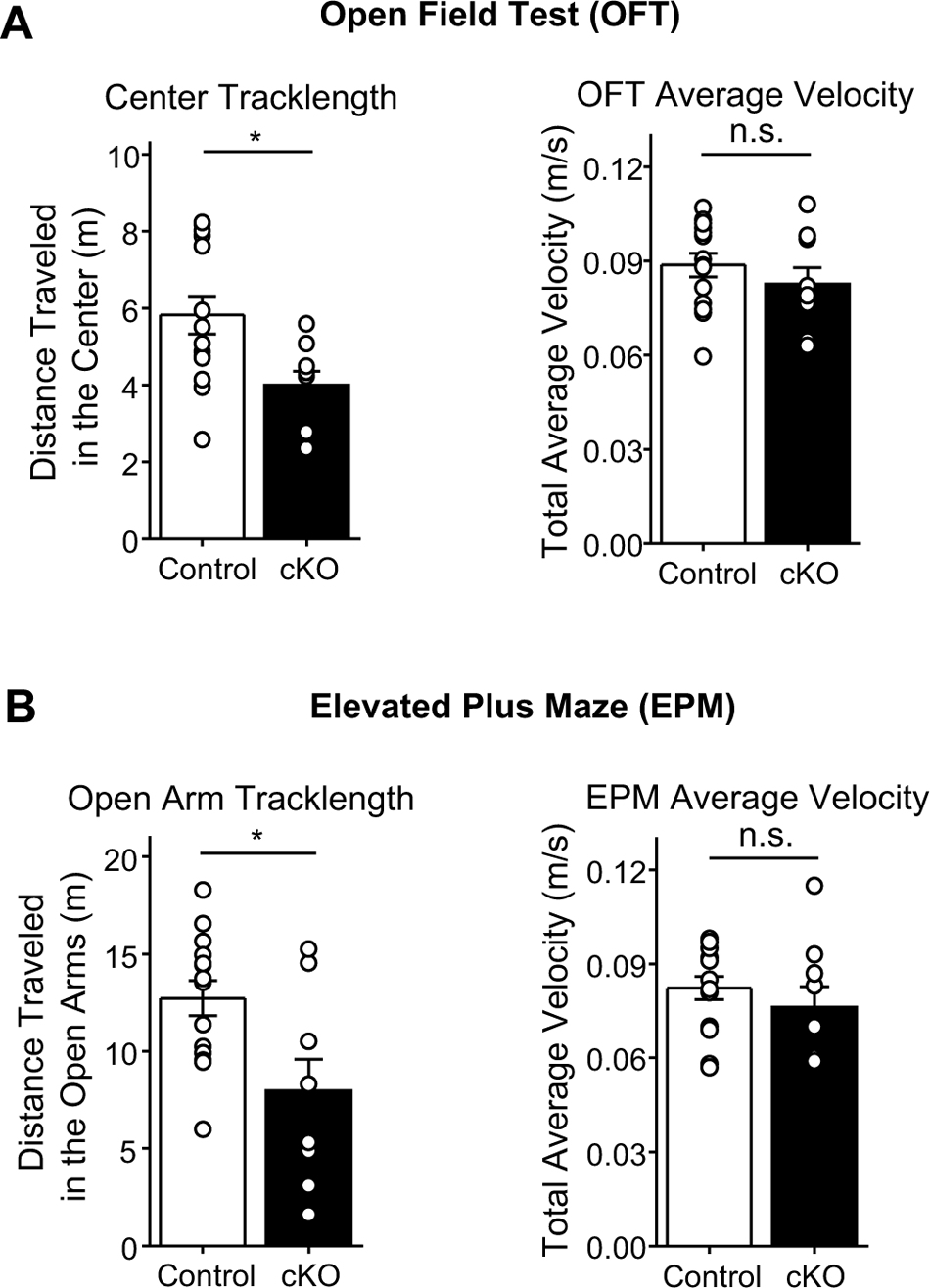
Additional features of MC *Drd2* cKO and Control mouse behavior in the Open Field Test and Elevated Plus Maze. **(A)** Additional features of mouse behavior in the Open Field Test (OFT). *Left*, MC *Drd2* cKO mice traveled a significantly shorter distance in the center of the OF as compared to Control mice (Control: 5.82 ± 0.49 m, N = 14; cKO: 3.99 ± 0.37 m, N = 9; Control vs cKO: p = 0.015, unpaired t-test). *Right*, There was no significant difference between Control and MC *Drd2* cKO mice in the average velocity of their travel in the entire arena area (Control: 0.0887 ± 0.0037 m/s, N = 14; cKO: 0.0827 ± 0.0051 m/s, N = 9; Control vs cKO: p = 0.34, unpaired t-test). **(B)** Additional features of mouse behavior in the Elevated Plus Maze (EPM). *Left*, MC *Drd2* cKO mice traveled a significantly shorter distance in the open arms of the EPM as compared to Control mice (Control: 12.7 ± 0.91 m, N = 14; cKO: 7.97 + 1.6, N = 9; Control vs cKO: p = 0.012, unpaired t-test). *Right*, There was no significant difference between Control and MC *Drd2* cKO mice in the average velocity of their travel in the entire maze (Control: 0.0823 + 0.0037 m/s, N = 14; cKO: 0.0762 + 0.0066 m/s, N = 9; Control vs cKO: p = 0.40, unpaired t-test). Data are represented as mean ± SEM. *p < 0.05, n.s. p > 0.2

**Figure S3.**
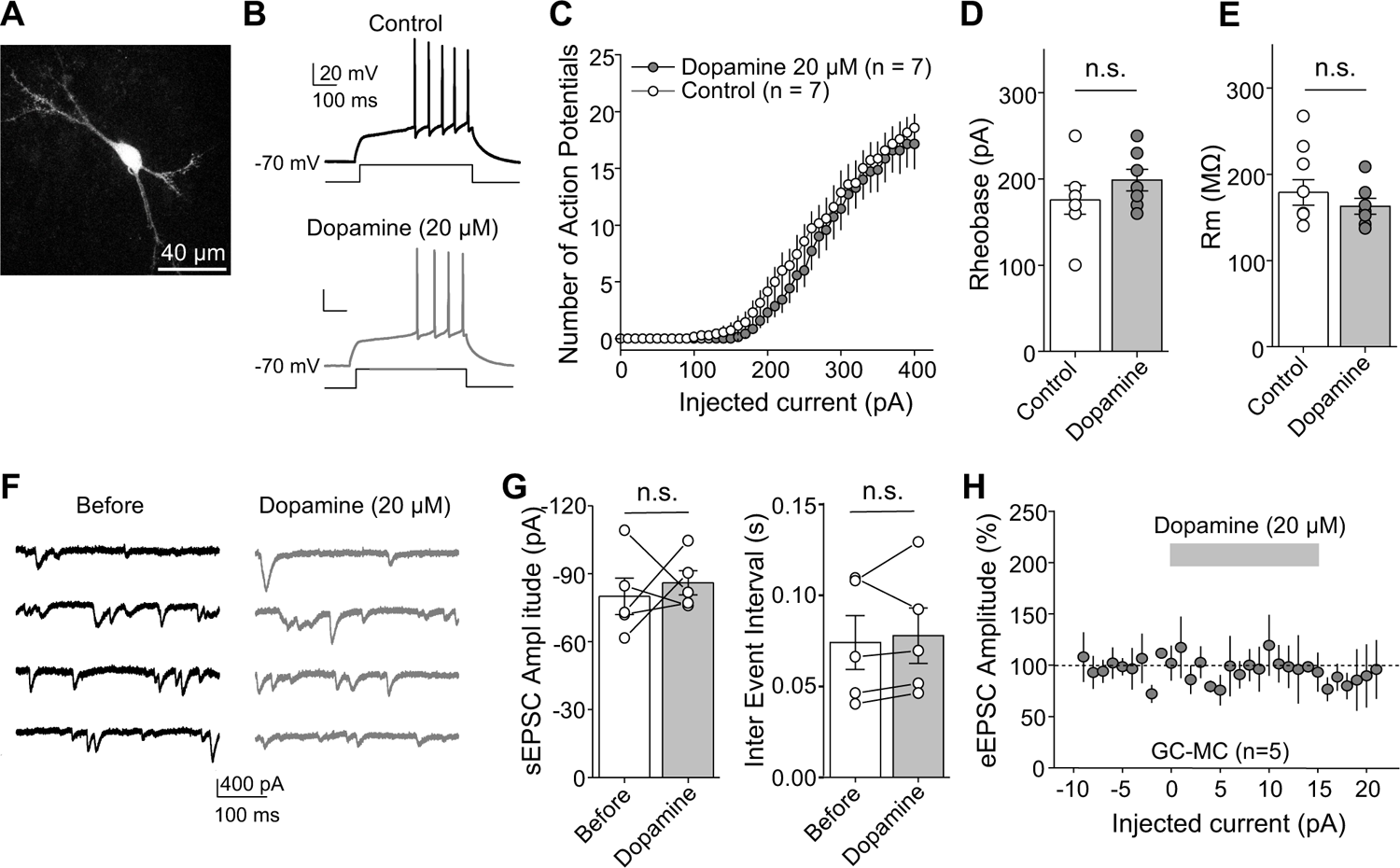
Dopamine bath application (20 µM) affected neither MC excitability nor excitatory input onto MCs. **(A)** Confocal image showing an example of a hilar MC filled with biocytin (white, Alexa Fluor 594-conjugated streptavidin). Note the presence of characteristic thorny excrescences on proximal MC dendrites. **(B-E)** In all recordings, synaptic transmission was blocked with NBQX (10 µM), D-APV (50 µM), picrotoxin (100 µM), and CGP55845 (3 µM). Dopamine (20 µM) or ACSF (control) was bath applied for 10 minutes prior to the start of and during MC recordings (V_holding_ = −70 mV, current clamp mode). Voltage responses were elicited using injections of 500 ms-long depolarizing current steps (from 0 pA to 400 pA, every 10 pA). Representative traces (B) and summary plot (C, D) showing that dopamine application did not significantly change the number of action potentials (C) or rheobase (D) of MCs (control: 175.7 ± 16.7%, n = 7; dopamine: 198.6 ± 12.4%, n = 7; control vs dopamine: p = 0.29, unpaired t-test). (E) Membrane resistance (Rm) was also not affected by dopamine application (control: 191.6 ± 17.9%, n = 7; dopamine: 162.8 ± 9.3%, n = 7; control vs dopamine: p = 0.19, unpaired t-test). **(F-H)** Spontaneous excitatory postsynaptic currents (sEPSCs) and GC-evoked EPSCs (eEPSCs) were recorded from MCs in the presence of picrotoxin (100 µM) and CGP55845 (3 µM). Single experiment representative traces (F) and summary plots (G) showing that dopamine application (20 µM) did not significantly change sEPSC amplitude (G, *left*, before vs dopamine: p = 0.62, n = 5, paired t-test) or inter event interval (G, *right*, before vs dopamine: p = 0.57, n = 5, paired t-test). (H) Dopamine application also did not affect the amplitude GC-evoked EPSCs (before vs dopamine: p = 0.98, paired t-test). Data are represented as mean ± SEM. n.s. p > 0.1

## References

1. D. M. Bannerman et al., Hippocampal synaptic plasticity, spatial memory and anxiety. Nature reviews. Neuroscience 15, 181–192 (2014).

2. T. Hainmueller, M. Bartos, Dentate gyrus circuits for encoding, retrieval and discrimination of episodic memories. Nature reviews. Neuroscience 21, 153–168 (2020).

3. B. A. Strange, M. P. Witter, E. S. Lein, E. I. Moser, Functional organization of the hippocampal longitudinal axis. Nature reviews. Neuroscience 15, 655–669 (2014).

4. H. E. Scharfman, The enigmatic mossy cell of the dentate gyrus. Nature reviews. Neuroscience 17, 562–575 (2016).

5. A. D. Bui et al., Dentate gyrus mossy cells control spontaneous convulsive seizures and spatial memory. Science (New York, N.Y.) 359, 787–790 (2018).

6. X. Li et al., A circuit of mossy cells controls the efficacy of memory retrieval by Gria2I inhibition of Gria2. Cell Rep 34, 108741 (2021).

7. S. Li et al., Alzheimer-like tau accumulation in dentate gyrus mossy cells induces spatial cognitive deficits by disrupting multiple memory-related signaling and inhibiting local neural circuit. Aging Cell 21, e13600 (2022).

8. J. J. Botterill, et al., Bidirectional Regulation of Cognitive and Anxiety-like Behaviors by Dentate Gyrus Mossy Cells in Male and Female Mice. J Neurosci 41, 2475-2495 (2021).

9. S. Jinde et al., Hilar mossy cell degeneration causes transient dentate granule cell hyperexcitability and impaired pattern separation. Neuron 76, 1189–1200 (2012).

10. K. Y. Wang et al., Elevation of hilar mossy cell activity suppresses hippocampal excitability and avoidance behavior. Cell Rep 36, 109702 (2021).

11. J. J. Botterill et al., An Excitatory and Epileptogenic Effect of Dentate Gyrus Mossy Cells in a Mouse Model of Epilepsy. Cell Rep 29, 2875–2889.e2876 (2019).

12. K. Nasrallah et al., Seizure-induced strengthening of a recurrent excitatory circuit in the dentate gyrus is proconvulsant. Proc Natl Acad Sci U S A 119, e2201151119 (2022).

13. P. S. Buckmaster, H. J. Wenzel, D. D. Kunkel, P. A. Schwartzkroin, Axon arbors and synaptic connections of hippocampal mossy cells in the rat in vivo. The Journal of comparative neurology 366, 271–292 (1996).

14. D. G. Amaral, H. E. Scharfman, P. Lavenex, The dentate gyrus: fundamental neuroanatomical organization (dentate gyrus for dummies). Prog Brain Res 163, 3–22 (2007).

15. G. Gangarossa et al., Characterization of dopamine D1 and D2 receptor-expressing neurons in the mouse hippocampus. Hippocampus 22, 2199–2207 (2012).

16. E. Puighermanal et al., drd2-cre:ribotag mouse line unravels the possible diversity of dopamine d2 receptor-expressing cells of the dorsal mouse hippocampus. Hippocampus 25, 858–875 (2015).

17. B. Missale, S. R. Nash, S. W. Robinson, M. Jaber, M. G. Caron, Dopamine receptors: from structure to function. Physiol Rev 78, 189–225 (1998).

18. S. K. Goldsmith, J. N. Joyce, Dopamine D2 receptor expression in hippocampus and parahippocampal cortex of rat, cat, and human in relation to tyrosine hydroxylase- immunoreactive fibers. Hippocampus 4, 354–373 (1994).

19. Y. Senzai, G. Buzsaki, Physiological Properties and Behavioral Correlates of Hippocampal Granule Cells and Mossy Cells. Neuron 93, 691–704.e695 (2017).

20. C. Jung et al., Dentate granule and mossy cells exhibit distinct spatiotemporal responses to local change in a one-dimensional landscape of visual-tactile cues. Sci Rep 9, 9545 (2019).

21. J. J. Botterill, K. J. Gerencer, K. Y. Vinod, D. Alcantara-Gonzalez, H. E. Scharfman, Dorsal and ventral mossy cells differ in their axonal projections throughout the dentate gyrus of the mouse hippocampus. Hippocampus 31, 522–539 (2021).

22. G. Etter, W. Krezel, Dopamine D2 receptor controls hilar mossy cells excitability. Hippocampus 24, 725–732 (2014).

23. D. G. McNamara, A. Tejero-Cantero, S. Trouche, N. Campo-Urriza, D. Dupret, Dopaminergic neurons promote hippocampal reactivation and spatial memory persistence. Nature neuroscience 17, 1658–1660 (2014).

24. K. A. Kempadoo, E. V. Mosharov, S. J. Choi, D. Sulzer, E. R. Kandel, Dopamine release from the locus coeruleus to the dorsal hippocampus promotes spatial learning and memory. Proc Natl Acad Sci U S A 113, 14835-14840 (2016).

25. T. Takeuchi et al., Locus coeruleus and dopaminergic consolidation of everyday memory. Nature 537, 357–362 (2016).

26. M. Piri, E. Ayazi, M. R. Zarrindast, Involvement of the dorsal hippocampal dopamine D2 receptors in histamine-induced anxiogenic-like effects in mice. Neurosci Lett 550, 139–144 (2013).

27. Y. Bozzi, D. Vallone, E. Borrelli, Neuroprotective role of dopamine against hippocampal cell death. J Neurosci 20, 8643–8649 (2000).

28. Y. Bozzi, E. Borrelli, The role of dopamine signaling in epileptogenesis. Frontiers in cellular neuroscience 7, 157 (2013).

29. M. Dunleavy, G. Provenzano, D. C. Henshall, Y. Bozzi, Kainic acid-induced seizures modulate Akt (SER473) phosphorylation in the hippocampus of dopamine D2 receptor knockout mice. J Mol Neurosci 49, 202–210 (2013).

30. A. Wilkerson, E. D. Levin, Ventral hippocampal dopamine D1 and D2 systems and spatial working memory in rats. Neuroscience 89, 743–749 (1999).

31. K. Yang, et al., Dopamine receptor activity participates in hippocampal synaptic plasticity associated with novel object recognition. Eur J Neurosci 45, 138–146 (2017).

32. H. Du, et al., Dopaminergic inputs in the dentate gyrus direct the choice of memory encoding. Proc Natl Acad Sci U S A 113, E5501-5510 (2016).

33. E. A. Petter, et al., Elucidating a locus coeruleus-dentate gyrus dopamine pathway for operant reinforcement. Elife 12 (2023).

34. A. J. Duszkiewicz, C. G. McNamara, T. Takeuchi, L. Genzel, Novelty and Dopaminergic Modulation of Memory Persistence: A Tale of Two Systems. Trends in neurosciences 42, 102–114 (2019).

35. A. Sík, N. Hájos, A. Gulácsi, I. Mody, T. F. Freund, The absence of a major Ca2+ signaling pathway in GABAergic neurons of the hippocampus. Proc Natl Acad Sci U S A 95, 3245-3250 (1998).

36. J. E. Zachry et al., Sex differences in dopamine release regulation in the striatum. Neuropsychopharmacology 46, 491–499 (2021).

37. O. O. F. Williams, M. Coppolino, S. R. George, M. L. Perreault, Sex Differences in Dopamine Receptors and Relevance to Neuropsychiatric Disorders. Brain Sci 11 (2021).

38. A. Vogel-Ciernia, M. A. Wood, Examining object location and object recognition memory in mice. Curr Protoc Neurosci 69, 8.31.31–17 (2014).

39. N. J. Broadbent, L. R. Squire, R. E. Clark, Spatial memory, recognition memory, and the hippocampus. Proc Natl Acad Sci U S A 101, 14515-14520 (2004).

40. M. R. Hunsaker, J. S. Rosenberg, R. P. Kesner, The role of the dentate gyrus, CA3a,b, and CA3c for detecting spatial and environmental novelty. Hippocampus 18, 1064-1073 (2008).

41. F. Fredes, R. Shigemoto, The role of hippocampal mossy cells in novelty detection. Neurobiol Learn Mem 183, 107486 (2021).

42. P. Simon, R. Dupuis, J. Costentin, Thigmotaxis as an index of anxiety in mice. Influence of dopaminergic transmissions. Behav Brain Res 61, 59–64 (1994).

43. A. A. Walf, C. A. Frye, The use of the elevated plus maze as an assay of anxiety-related behavior in rodents. Nat Protoc 2, 322–328 (2007).

44. M. Levesque, M. Avoli, The kainic acid model of temporal lobe epilepsy. Neurosci Biobehav Rev 37, 2887–2899 (2013).

45. B. G. Robinson, et al., Desensitized D2 autoreceptors are resistant to trafficking. Sci Rep 7, 4379 (2017).

46. J. J. Lebowitz et al., Subcellular localization of D2 receptors in the murine substantia nigra. Brain Struct Funct 227, 925–941 (2022).

47. E. S. Calipari, M. J. Ferris, Amphetamine mechanisms and actions at the dopamine terminal revisited. J Neurosci 33, 8923–8925 (2013).

48. T. Tsetsenis, J. I. Broussard, J. A. Dani, Dopaminergic regulation of hippocampal plasticity, learning, and memory. Frontiers in Behavioral Neuroscience 16 (2023).

49. J. Rocchetti et al., Presynaptic D2 dopamine receptors control long-term depression expression and memory processes in the temporal hippocampus. Biol Psychiatry 77, 513–525 (2015).

50. Y. Mu, C. Zhao, F. H. Gage, Dopaminergic modulation of cortical inputs during maturation of adult-born dentate granule cells. J Neurosci 31, 4113-4123 (2011).

51. Y. Hashimotodani et al., LTP at Hilar Mossy Cell-Dentate Granule Cell Synapses Modulates Dentate Gyrus Output by Increasing Excitation/Inhibition Balance. Neuron 95, 928–943.e923 (2017).

52. K. R. Jensen, C. Berthoux, K. Nasrallah, P. E. Castillo, Multiple cannabinoid signaling cascades powerfully suppress recurrent excitation in the hippocampus. Proc Natl Acad Sci U S A 118 (2021).

53. C. M. Lovinger et al., Local modulation by presynaptic receptors controls neuronal communication and behaviour. Nature reviews. Neuroscience 23, 191–203 (2022).

54. J. K. Leutgeb, S. Leutgeb, M. B. Moser, E. I. Moser, Pattern separation in the dentate gyrus and CA3 of the hippocampus. *Science (New York*, N.Y.) 315, 961–966 (2007).

55. T. J. McHugh et al., Dentate gyrus NMDA receptors mediate rapid pattern separation in the hippocampal network. *Science (New York*, N.Y.) 317, 94–99 (2007).

56. A. Treves, E. T. Rolls, Computational analysis of the role of the hippocampus in memory. Hippocampus 4, 374–391 (1994).

57. R. P. Kesner, E. T. Rolls, A computational theory of hippocampal function, and tests of the theory: new developments. Neurosci Biobehav Rev 48, 92–147 (2015).

58. G. D. Bartoszyk, Anxiolytic effects of dopamine receptor ligands: I. Involvement of dopamine autoreceptors. Life Sci 62, 649–663 (1998).

59. K. Nasrallah, et al., Retrograde adenosine/A 2A receptor signaling mediates presynaptic hippocampal LTP and facilitates epileptic seizures. *bioRxiv* 10.1101/2021.10.07.463512, 2021.2010.2007.463512 (2021).

60. B. R. Butler, G. L. Westbrook, E. Schnell, Adaptive Mossy Cell Circuit Plasticity after Status Epilepticus. J Neurosci 42, 3025-3036 (2022).

## SI Appendix References

1. V. Brust, P. M. Schindler, L. Lewejohann, Lifetime development of behavioural phenotype in the house mouse (Mus musculus). Front Zool 12 Suppl 1, S17 (2015).

2. J. J. Botterill, et al., Bidirectional Regulation of Cognitive and Anxiety-like Behaviors by Dentate Gyrus Mossy Cells in Male and Female Mice. J Neurosci 41, 2475-2495 (2021).

3. A. Vogel-Ciernia, M. A. Wood, Examining object location and object recognition memory in mice. Curr Protoc Neurosci 69, 8.31.31–17 (2014).

4. R. J. Racine, Modification of seizure activity by electrical stimulation. II. Motor seizure. Electroencephalogr Clin Neurophysiol 32, 281–294 (1972).

5. J. J. Lebowitz et al., Subcellular localization of D2 receptors in the murine substantia nigra. Brain Struct Funct 227, 925–941 (2022).

6. B. G. Robinson, et al., Desensitized D2 autoreceptors are resistant to trafficking. Sci Rep 7, 4379 (2017).

7. C. Schmidt-Hieber, P. Jonas, J. Bischofberger, Enhanced synaptic plasticity in newly generated granule cells of the adult hippocampus. Nature 429, 184–187 (2004).

8. P. Larimer, B. W. Strowbridge, Nonrandom local circuits in the dentate gyrus. J Neurosci 28, 12212-12223 (2008).

9. T. P. Hedrick, et al., Excitatory Synaptic Input to Hilar Mossy Cells under Basal and Hyperexcitable Conditions. eNeuro 4 (2017).

10. M. Lysetskiy, C. Földy, I. Soltesz, Long- and short-term plasticity at mossy fiber synapses on mossy cells in the rat dentate gyrus. Hippocampus 15, 691–696 (2005).

11. D. G. Amaral, A Golgi study of cell types in the hilar region of the hippocampus in the rat. The Journal of comparative neurology 182, 851–914 (1978).

12. E. G. Amaral, H. E. Scharfman, P. Lavenex, The dentate gyrus: fundamental neuroanatomical organization (dentate gyrus for dummies). Prog Brain Res 163, 3–22 (2007).

13. A. T. Trinh, M. Girardi-Schappo, J. C. Béïque, A. Longtin, L. Maler, Dentate gyrus mossy cells exhibit sparse coding via adaptive spike threshold dynamics. bioRxiv 10.1101/2022.03.07.483263, 2022.2003.2007.483263 (2022).

